# The cyclin-G associated kinase (GAK) is a novel mitotic kinase and therapeutic target in diffuse large B-cell lymphoma

**DOI:** 10.1101/2024.10.21.619489

**Authors:** Olivia B. Farag, Bassel Awada, Abdessamad A. Youssfi, Paola Manara, Anna Kingham, Preet Kumar, Lingxiao Li, Brian B. Silver, Toral Shastri, Santiago Vilar, Austin D. Newsam, Marianna Lekakis, Diego E. Hernandez Parets, Pierre A. Younes, Dhanvantri Chahar, Artavazd Arumov, Caroline A. Coughlin, Alicia Bilbao-Martinez, Mahesh Gokara, Francesco Maura, Stephan C. Schürer, Yangbo Feng, Vance Lemmon, Daniel Bilbao, Hassan Al-Ali, Jonathan H. Schatz

## Abstract

New drug targets are needed for diffuse large B-cell lymphoma (DLBCL), the most common lymphoma subtype, to enable enable development better treatments for patients not cured by standard care. We conducted a phenotypic screen of kinase inhibitors and identified the cyclin G-associated kinase (GAK) as a tumor-selective target. Though GAK is previously described primarily as a participant in membrane trafficking, we found its kinase activity is a key mitotic regulator in DLBCL. Inhibition caused G2/M-phase arrest, chromosome misalignment, and spindle distortion, effects absent in non-malignant controls. Transcriptomics data from clinical samples showed increased *GAK* expression associates with *RB1* deficiency in DLBCL cases, suggesting dependency on GAK linked to retinoblastoma associated protein (RB) loss of function, a common DLBCL driver. RB-deficient DLBCL cells treated with a selective GAK tool compound showed complete arrest at G2/M, pronounced distortion of mitotic spindles, and widespread chromosomal damage. High-content live-cell imaging revealed onset of mitotic catastrophe in response to GAK inhibition, which was more rapid and severe in isogenic cells with *RB1* deletion. No GAK-selective inhibitors suitable for clinical development are currently available, but several drugs approved or under development inhibit GAK activity even more potently than thier intended clinical targets. For instance, OTS167, developed against MELK for use in solid tumors, has particularly potent anti-GAK potency and has achieved single-agent tumor-burden reduction in vivo against a DLBCL patient-derived xenograft. GAK is therefore a novel mitotic kinase in DLBCL, linked to the common, undruggable RB loss of function biomarker, and suitable for rapid clinical translation through drug repurposing.

**Significance:** We identify cyclin-G associated kinase (GAK) as a novel therapeutic vulnerability in diffuse large B-cell lymphoma. Clinical kinase inhibitors with GAK activity create an opportunity for rapid therapeutic translation through drug repurposing.

## Introduction

Diffuse large B-cell lymphoma (DLBCL) is among the most frequently diagnosed hematologic malignancies, comprising a clinically and molecularly heterogeneous group of sub-entities.^1^ Gene-expression profiling two decades ago divided DLBCL into germinal center B-cell (GCB) and activated B-cell (ABC) cell-of-origin (COO) subtypes, with the latter carrying worse prognosis.^2,3^ Subsequent genomics studies defined 5-7 molecular subgroups based on varying frequency of recurrent alterations.^4^ Despite greatly increased comprehension of disease biology, however, there has been little progress developing interventions directed at specific molecular subtypes. R-CHOP (rituximab, cyclophosphamide, doxorubicin, vincristine, and prednisone) chemoimmunotherapy remains standard frontline treatment regardless of molecular characteristics.^5^ A recent substitution of vincristine with the anti-CD79B antibody-drug conjugate polatuzumab vedotin (pola-R-CHP) introduced a new alternative with improved progression-free survival, but did not impact overall survival.^6^ Frontline treatment choices remain largely uninformed by the prognostic significance of molecular biomarkers, and at least 30-40% of patients will not achieve cure.^7^ CD19-directed chimeric antigen receptor (CAR)-T cells are potentially curative new salvage options.^8^ However, these resource-intensive therapies are only available at specialized centers, and a third or less of patients achieve cure from this approach.^9^

Targeted therapies such as Bruton’s tyrosine kinase (BTK) inhibitors have revolutionized care of other B-lymphomas but strikingly have found little role in the management of DLBCL.^10^ ViPOR (venetoclax, ibrutinib, prednisone, obitutuxumab, lenalidomide) showed promising response rates in heavily pretreated patients and is the first active DLBCL regimen with a kinase inhibitor as well as the first to exclude traditional chemotherapy agents completely.^11^ However, long-term assessment of response duration remains unclear. Currently, salvage chemoimmunotherapy, requiring stem cell transplant consolidation in second-line settings, and CAR-T therapy remain the only potentially curative treatment paradigms for relapsed or refractory (rel/ref) DLBCL. Most of these patients unfortunately still die from complications of disease progression.^12^

Utilizing a suite of phenotypic screens with DLBCL cell lines and healthy cells, complemented by machine learning-based target deconvolution, we identified the cyclin-G associated kinase (GAK) as a novel, cancer-selective target in both GCB and ABC DLBCL. GAK has been previously characterized as a participant in clathrin-mediated endocytosis (CME) and vesicular trafficking, but here we find critical roles for its kinase function in successful mitosis.^13–16^ This study is the first to establish proof-of-principle for GAK as a novel and tractable therapeutic target in DLBCL or other malignancies. Critically, we overcame barriers to in vivo evaluation to establish proof-of-principle for GAK as a novel and therapeutic target. These findings provide a rationale for therapeutic targeting of GAK and highlight opportunities for rapid clinical translation through repurposing of existing kinase inhibitors with GAK activity.

## Methods

### Cell Lines and Reagents

293T and U2OS cells were obtained from the American Type Culture Collection (ATCC). U2OS *RB1*-/- CRISPR-modified cells were gifted to us by Dr. Fred Dick at Western University, Ontario. 293T, U2OS, A549, HeLa, and U87 cell lines were cultured in DMEM with 10% FBS, 1% penicillin-streptomycin and Plasmocin® (InvivoGen, Cat# ant-mpp). Karpas 422, SU-DHL-6, Toledo, WSU-DL-2, RIVA, SU-DHL-4, U2932, HBL1, TMD8, PBMC, SU-DHL-2, Farage, Ly19, DB, Ly8, HL60, K562, MOLT-4, RS4;11, MAC2A, SUPM2, DHL-1, Karpas 299, Raji, H2228, BJAB, and HCC15 cell lines were obtained from ATCC and cultured in RPMI plus 10% FBS, 1% penicillin-streptomycin, and Plasmocin®. Ly1 was obtained from the Deutsche Sammlung von Mikroorganismen und Zellkulturen (DSMZ) and cultured in IMDM plus 10% FBS, 1% penicillin-streptomycin, and Plasmocin®. Cells were maintained in an incubator at 37*°*C with 5% CO_2_ and all cell lines were verified by STR finger printing and tested for mycoplasma. Sources of compounds in cell-based experiments are in **Table S1**.

### Phenotypic Screen

The Published Kinase Inhibitor Set (PKIS-I) library compounds were screened against 10 DLBCL cell lines (TMD8, HBL1, U2932, SU-DHL4, SU-DHL10, RIVA, Toledo, SU-DHL2, SU-DHL6, Karpas422) and PBMCs. The drug sensitivity screen was performed as previously described.^17^ Compounds were plated into white, clear-bottom 384-well plates at 5 different concentrations, using 10-fold serial dilutions to cover a 10,000-fold concentration range (1 nM to 10 μM). Liquid handling for compound preparation was carried out using a PerkinElmer Janus dual-head robot. The pre-plated compounds were dissolved in 5 μL of culture medium and incubated for 30 minutes at room temperature on an orbital shaker. 20 μL cell suspension was added to the treatment plates. After 72 hours of incubation, 2.5 μL of CellTiter-Blue viability reagent (Promega) was added to each well. The plates were incubated for an additional 2 hours at 37°C, and fluorescence was measured using a PHERAstar FS plate reader (BMG Labtech). The drug sensitivity score (DSS), based on the area under the dose-response curve, was calculated for each compound using the DSSR software package.^17^ The differential DSS (dDSS) was then determined for each compound in each cell line using the formula: dDSS = DSS (cell line) - DSS (PBMC).

### Target Deconvolution Analysis

Average dDSS values were calculated for each compound across all 10 DLBCL cell lines then used to stratify compounds into one of two classes: hits (dDSS ≥ 10) and non-hits (dDSS ≤ 2). Kinase activity profiles for the stratified compounds were acquired from previously published data.(Drewry, Wells et al. 2017) In total, 375 compounds and 386 non-mutant kinases were included in the analysis. Target deconvolution was performed using previously described computational methods.^17^ The top 10% scoring kinase groups were prioritized for orthogonal validation with RNAi.

### siRNA Validation

Dharmacon Acell siRNAs from Horizon were used for this study. The siRNA stock solution was prepared by diluting 5x siRNA Buffer (Cat #B-002000-UB-100) to a 1x concentration using sterile RNase-free water. The siRNA was then resuspended to a final concentration of 100 μM in 1x siRNA Buffer. The solution was gently mixed, incubated at 37 °C for 70 minutes, and transferred to 384 well plates. For each transfection, 500 DLBCL cells were suspended in Accell siRNA Delivery Media (Cat #B-005000) and combined with the siRNA solution (initially 100μM) to achieve a final siRNA concentration of 1 μM. The plates were then incubated at 37°C with 5% CO2 for 72 hours. The viability of cells after siRNA treatment was assessed with CellTiter-Glo (Promega). For the initial validation, siRNAs were tested in triplicate and effects were calculated as robust z-scores. The differential robust z-scores for each siRNA target in each cell line were calculated according to the formula d_robust_z-score = robust z-score (cell line) – robust z-score (PBMCs) **(Table S3)**. For follow-up validation, the siRNA treatments were conducted in four technical replicates, and the effects were converted to percent viability compared to those observed in the parental cell lines.

### Cell Viability Assay

Cells were seeded in 96-well plates at a density of 3,000-5,000 cells per well. The cells were treated with compounds using 1:2 serial dilution and incubated for 48 or 72 hours at 37 °C with 5% CO2. For adherent cell lines, cells were seeded and allowed to adhere overnight before treatment. Viability was assessed using luminescence measurements with CellTiter-Glo (Promega), and percent viability was calculated by normalizing readings to vehicle (DMSO) treatment. A nonlinear fit regression analysis was performed using GraphPad Prism to determine IC_50_. For synergy assays, cells were incubated with drug combinations for 72 hours. Viability data were analyzed using the SynergyFinder tool to calculate Bliss Synergy Score.

### Growth Inhibition

Suspension cell lines were seeded in 96-well plates at a density of 3,000-5,000 cells per well. The cells were treated with the compounds using 1:2 serial dilution and incubated at 37 °C with 5% CO2. For adherent cell lines, cells were seeded and allowed to adhere overnight before treatment. After 48 hours, the treatment was refreshed by adding additional media and compound to wells. Growth inhibition was assessed at 96 hours using luminescence measurements with the CellTiter-Glo assay (Promega), and percent growth inhibition was calculated normalized to vehicle (DMSO) treatment. A nonlinear fit regression analysis was performed using GraphPad Prism to determine GI_50_.

### Western Blot

For sample preparation, cells were pelleted by centrifugation at 1100 rpm for 5 minutes. The supernatant was removed, and the cells were washed with cold PBS. Cell lysis was performed using RIPA buffer containing 1x Triton X (Thermo Fisher Scientific) along with a protease/phosphatase inhibitor cocktail (Thermo Fisher Scientific). The lysate was then centrifuged, and the supernatant was collected. Total protein concentration was quantified using the Bradford reagent (Bio-Rad). Proteins were separated on Bolt™ 4–12% Bis-Tris Plus gels (Thermo Fisher Scientific) using the Mini Gel Tank system (Thermo Fisher Scientific). Following electrophoresis, proteins were transferred to 0.2 μm Immuno-Blot PVDF membranes (Bio-Rad) at 26V for one hour using the Bolt™ Mini Blot Module (Thermo Fisher Scientific), following the manufacturer’s instructions. Membranes were blocked with 5% milk and probed with primary antibodies **(Table S2)** followed by host-specific secondary antibodies. Chemiluminescent signals were measured using a digital chemiluminescent imaging system (LiCor).

### BCR Stimulation and Internalization Assays

Cells were seeded at a density of 1 × 10⁶ cells/mL and treated with DMSO vehicle control, SGC-GAK-1 (1 µM), or dynasore (100 µM) for 2 hours prior to stimulation. Drug treatment was maintained throughout stimulation and subsequent incubation periods.For BCR activation western blot, cells were stimulated by incubation with goat anti-human IgM (10 µg/mL) for 30 minutes at 4°C to allow surface binding. Cells were then resuspended in pre-warmed media containing the respective drug and incubated at 37°C for 20 minutes. Stimulation was terminated by transferring samples to ice and washing with cold PBS. Cells were lysed, and protein extracts were subjected to immunoblot analysis. For FACS-based BCR internalization assay, following surface IgM binding and washing, cells were resuspended in pre-warmed media containing drug and incubated at 37°C for 10, 20, 30, or 60 minutes. Internalization was halted by transferring cells to ice and washing with cold PBS. Cells were incubated with FITC-conjugated donkey anti-goat IgG (H+L) secondary antibody (1:2000) for 45 minutes on ice to detect remaining surface-bound IgM. Flow cytometric analysis was performed, and surface IgM levels were quantified as the mean fluorescence intensity (MFI) of Alexa Fluor 488. MFI values were normalized to the 0-minute time point to assess relative BCR internalization over time.

### Transferrin Uptake

Cells were seeded at a density of 1 × 10⁶ cells/mL and treated with DMSO vehicle control, SGC-GAK-1 (1 µM), or dynasore (100 µM) for 2 hours. To reduce endogenous transferrin receptor occupancy, cells were serum-starved in serum-free media containing the respective drug treatment for 30 minutes at 37°C. Cells were incubated on ice for 10 minutes to halt endocytosis. CF®488A-conjugated human transferrin (5 µg/mL) was added in ice-cold serum-free media (+ drug), and cells were incubated for 15 minutes at 4°C to allow surface binding. Unbound transferrin was removed by washing with ice-cold PBS supplemented with 1% FBS. Cells were resuspended in pre-warmed complete media containing the respective drug and incubated at 37°C for 5 minutes to allow transferrin uptake. Uptake was terminated by transferring cells to ice followed by washing with ice-cold PBS. Transferrin uptake was quantified by flow cytometry. MFI of CF®488A of was calculated, and values were normalized to the DMSO control or non-targeting (NT) siRNA control as indicated.

### RNA Sequencing

U2932 or Ly1 cells were treated in triplicate with either 1:1000 DMSO (vehicle control) or SGC-GAK-1 at a concentration of 1 µM for 12 and 24 hours. RNA extraction and sample processing was conducted at the John P. Hussman Institute for Human Genomics core at the University of Miami. Total RNA was extracted using the Qiagen Symphony system and quantified with Qubit. Total RNA was prepped with the Nugen Universal Plus mRNA-Seq (M01442 v2) using 500ng via Qubit following poly-A selection and 14 cycles of PCR amplification. Libraries were sequenced on an Illumina Novaseq X platform, generating approximately 30 million PE150 reads per sample. Raw sequencing reads were quality-checked using FastQC, and reads were aligned against the GRCh38 human reference genome using the STAR aligner. Differential gene expression analysis was performed using edgeR and limma, with a cut off of log-2 fold change > 0.5 and adjusted p-value < 0.05. Unsupervised clustering of differentially expressed genes was used to assess treatment-specific gene expression profiles. Gene-set enrichment analysis (GSEA) was conducted using the FGSEA package, with a false discovery rate (FDR) q value threshold set at <0.25.

### FACS-based Cell Cycle Analysis

Cells were stained with Vybrant™ DyeCycle™ Orange Stain (Thermo Fisher) according to the manufacturer’s protocol. Cells were suspended in complete medium at a concentration of 1 × 10^6 cells/mL. Vybrant DyeCycle Orange was added to each sample at a final concentration of 10 μM, and the cells were incubated at 37°C for 30 minutes, protected from light. After staining, samples were analyzed using a flow cytometer with 488 nm excitation and orange emission. Data analysis was performed using FlowJo software to assess cell cycle distribution.

### Immunofluorescent Confocal Imaging

Adherent cells (U2OS): 250k cells per well were seeded on glass coverslips placed in 12-well plates. Cells were allowed to adhere overnight, followed by treatment with SGC-GAK-1 or DMSO (vehicle control). Following treatment, cells were fixed and stained directly on the coverslips.

Suspension cells (U2932, RIVA, TMD8, PBMC): 1 million cells per well were seeded in 6-well plates and treated with SGC-GAK-1 or DMSO (vehicle control). Following treatment, cells were fixed and washed by centrifugation and resuspension. Resuspended, fixed cells were plated on poly-lysine coated coverslips and incubated at room temperature for 20 minutes to allow cells to adhere to the coverslips prior to staining. All cells were fixed with 4% paraformaldehyde for 10 minutes at room temperature, then washed three times with PBS (5 minutes per wash). All incubations were performed light-protected at room temperature. Cells were blocked with blocking buffer (Cell Signaling #12411) for 30 minutes. The blocking solution was removed, and cells were incubated with primary antibody (anti-alpha tubulin, Thermo Fisher, #14-4502-82) diluted in blocking buffer for 1 hour. After washing three times with PBS (5 minutes each), cells were incubated with secondary antibody (Goat anti-Mouse Alexa Fluor™ 488, Thermo Fisher, #A-10680) and DAPI (Thermo Fisher, #62248) diluted in blocking buffer, for 30 minutes at room temperature. After washing with PBS and Milli-Q water, the slides were dried in the dark at room temperature. Coverslips were mounted with ProLong™ Diamond Antifade Mountant (Thermo Fisher #P36965) and sealed with nail polish. Imaging was performed on a Leica confocal microscope using LAS X software. DAPI quantification for cell cycle analysis was conducted using LAS X software, and statistical significance was assessed with unpaired t-tests.

### Live Cell Painting Assay

U2OS WT and RB1-/- cells were seeded in 96-well plates (Corning, #3904) and allowed to adhere overnight at 37°C with 5% CO₂. WT cells were seeded at a density of 4000/well, while RB1-/- cells were seeded at a density of 2000/well to accommodate their higher proliferation rate. Cells were stained using the PhenoVue Live Cell Painting Kit (Revvity, #PLIVCP14) according to the manufacturer’s recommendations (PhenoVue Hoechst 33342 nuclear stain, PhenoVue 488 vesicle stain, PhenoVue 555 mitochondrial stain, PhenoVue 647 tubulin stain). Briefly, a dye mix was prepared in culture medium, added to the cells, and incubated for 4 hours at 37°C. Following staining, cells were treated with vehicle (0.1% DMSO) or SGC-GAK-1 (2 or 10 µM), with three technical replicates per condition for each cell line. Plates were imaged at 2-hour intervals in an Opera Phenix Plus High-Content Screening System (Perkin Elmer) equipped with an Environmental Control Unit. Images were acquired using a 40X objective in confocal mode. A total of 81 fields (9×9) were acquired from the center of each well.

### Cell Painting Assay Image and Data Analyses

Images were analyzed using the Cell Painting block in Harmony software. Feature values were computed as the median per well. Cell counts were determined for each well and normalized to the average of vehicle-treated wells at 48 hours for each cell line. 4,710 morphological features were extracted from Cell Painting images and filtered by excluding biologically nonsensical channel-compartment combinations: vesicle and mitochondria in the nucleus compartment, DNA in the membrane region, and redundant whole-cell composite features, resulting in 3,080 biologically curated features. Hoechst signal in the cytoplasm and ring regions was retained to capture micronuclei and mitotic chromosome morphology. Alexa 647 signal in the nucleus region was retained to capture spindle morphology during mitosis. Feature values were standardized to a zero mean and unit variance across samples within each cell line before dimensionality reduction. Principal component analysis (PCA) was performed, and components exceeding 90% cumulative explained variance were used to compute pairwise Euclidean distances between samples in PC space. Pairwise distances were used as input to metric multidimensional scaling (MDS), producing a 2D embedding in which inter-point distances reflect morphological similarity. Embedding quality was assessed using Kruskal stress-1 (<0.2 considered good). For each morphological feature, effect size was quantified using Cohen’s d, calculated as the difference between each treatment condition mean and the WT vehicle control mean, divided by the pooled standard deviation across replicates. All analyses were performed in MATLAB using the Statistics and Machine Learning Toolbox. From the 3,080 biologically curated features, a subset was selected for detailed biological interpretation. Features were required to exceed |Cohen’s d| > 5 in at least one treatment condition. Selection prioritized non-redundant biological readouts across four phenotypic domains informed by the known cellular functions of GAK and the established consequences of RB1 loss. When multiple features captured the same readout across different texture filters, representatives were retained based on effect size magnitude and complementarity of the texture preprocessing (e.g., spot-detection vs. spot-positive cell filtering). The within-RB1-/- subtraction isolates drug-induced changes from the constitutive RB1-/- phenotypic shift, enabling direct comparison of treatment response magnitude between genotypes.

### NanoBRET® Target Engagement (TE) Intracellular Kinase Assay

The NanoBRET® TE Intracellular Kinase Assay kit (Promega, #N2640) and [NanoLuc®-GAK Fusion Vector](https://www.promega.com/products/cell-signaling/kinase-target-engagement/nanoluc-gak-fusion-vector/) (Promega, NV142A) were used according to the manufacturer’s protocol. 293T cells were transfected with [NanoLuc®-GAK Fusion Vector](https://www.promega.com/products/cell-signaling/kinase-target-engagement/nanoluc-gak-fusion-vector/) using FuGENE® HD Transfection Reagent and seeded in white-bottom 96-well plates at a density of 2 × 10^5 cells/mL in Assay Medium. Cells were incubated for 20–30 hours post-transfection. After incubation, NanoBRET™ Tracer Reagent was added to the cells, followed by test compounds serially diluted in Opti-MEM I Reduced Serum Medium. The plates were incubated for 2 hours at 37°C and 5% CO2. Following incubation, 3X NanoBRET™ Substrate plus Inhibitor Solution was added, and BRET signals were measured using a luminometer at donor and acceptor wavelengths of 450nm and 610nm, respectively. The BRET ratio was calculated as the ratio of acceptor to donor emission values, and raw BRET units were converted to milliBRET units (mBU). A nonlinear fit regression analysis was performed using GraphPad Prism to determine Kd.

### *In vivo* Experiments

All mouse studies were performed under the approval of the University of Miami Institutional Animal Care and Use committee. *In vivo* experiments were conducted at the Cancer Modeling Shared Resource at the University of Miami. The mice were housed within a barrier facility. All equipment, food, or bedding exposed to the mice was autoclaved, irradiated (food), or chemically disinfected. NSG mice (6–8 weeks old) were used for our in vivo study. Mice were divided into two experimental groups: vehicle control (10 mice) and ABT + SGC-GAK-1 treatment (10 mice). Luciferase-engineered U2932 cells (1 × 10^6) were injected intravenously via the tail vein into each mouse, and treatment began one day following injection. Mice in the ABT + SGC-GAK-1 group were administered 50 mg/kg of ABT via oral gavage 2 hours prior to 10 mg/kg of SGC-GAK-1, also via oral gavage. Vehicle-treated mice received 10% DMSO, 40% PEG 300, 5% Tween 80, and 45% saline. Treatments were administered daily. Mice were monitored three times a week for tumor burden using IVIS imaging and overall health by weight measurements. Mice were anesthetized using 1–5% vaporized isoflurane in 100% oxygen during procedures. If mice exhibited more than 20% weight loss over two consecutive measurements or developed morbidity (e.g., lethargy, splenomegaly, or tumors >2 cm^3), they were euthanized using CO2 asphyxiation followed by cervical dislocation. Ex vivo analysis was performed on collected organs. For the OTS167 study, NSG mice were subcutaneously implanted with GCB DLBCL patient-derived xenograft (PDX) tumors, and treatment was initiated once tumors reached ≥100 mm³ (ultrasound imaging). Mice were randomized (n=10 per group) to receive either OTS167 at 10 mg/kg or vehicle control (0.5% methylcellulose) administered orally five times per week. Treatment continued until endpoint criteria were met (tumor volume ≥1,500 mm³, ≥20% body weight loss, or up to 90 days).

### Immunohistochemistry

These experiments were carried out at histology laboratory of the Cancer Modeling Shared Resource at the University of Miami. Liver and lung tissues were harvested from mice, fixed in 10% formalin, and embedded in paraffin. Sections (5 µm) were deparaffinized and rehydrated. All staining was performed, using the Leica Bond RX automated stainer (Leica Microsystems). Epitope retrieval was performed by heat-induced epitope retrieval of the formalin-fixed, paraffin-embedded tissue using citrate-based pH 6 solution (Bond ER1 solution) for 20 min at 95 °C. The tissues were first incubated with peroxide block buffer (Leica Microsystems) for 5 minutes, followed by blocking with 10% normal goat serum for 30 minutes. Tissue sections were incubated with primary antibody against Ki67. After washing, the slides were treated with a biotinylated secondary antibody and counterstained with hematoxylin. Slides were washed to remove non-specific binding and were dried, cover slipped, and visualized using a Leica microsystem (CMS GmbH).

### Quantification and Statistical Analysis

Experiments were performed in three independent replicates unless otherwise noted. Statistical analyses employed GraphPad Prism version 9. Unpaired student’s t test and ANOVA statistical significance: ns, p > 0.05; *p ≤ 0.05; **p ≤ 0.01; ***p ≤ 0.001 and ****p ≤ 0.0001).

## Results

### Phenotypic Screening Identifies GAK as a Novel Drug Target Candidate in DLBCL

To identify novel drug targets for DLBCL across subtypes, we designed a multistep discovery pipeline incorporating phenotypic drug screening, computational target deconvolution, and hit validation (**Fig. 1A**). First, we determined activity of 490 kinase inhibitors from the PKIS-I, PKIS-II, and KCGS libraries against 6 GCB- and 4 ABC-derived DLBCL cell lines, with peripheral blood mononuclear cells (PBMCs) as a non-malignant control **(Fig. 1B)**.^18,19^ For each inhibitor, we calculated a Drug Sensitivity Score (DSS) and differential DSS (dDSS), subtracting the DSS for each inhibitor in PBMCs from the inhibitor’s DSS of each cell line **(Fig. S1A, Table S3-4)**.^17^ To focus the target deconvolution analysis on selective killing of cancer cells, a dDSS threshold of ≥10 was used to identify initial hit compounds. Kinase activity profiles for the stratified compounds were acquired from previously published sources and integrated as previously described.^17^ We then deployed the idTRAX target deconvolution algorithm to identify kinases whose inhibition tracks with selective killing of cancer cells **(Table S5)**.^17^ The established lymphoma drug target aurora kinase A (AURKA) emerged as a high priority hit, validating the approach, while the cyclin-G associated kinase (GAK) emerged as a novel candidate. The Aurora kinases have been explored as therapeutic targets in cancer due to critical roles regulating multiple steps of the cell cycle but yielded poor efficacy within acceptable limits of toxicity in clinical trials.^20^ In DLBCL for example, Phase II evaluations of the AURKA inhibitor alisertib found limited efficacy, with objective response rate (ORR) of <20%, and high rates of grade 3+ hematologic toxicities which discouraged further clinical trials.^21^ We therefore focused on GAK, an understudied kinase and completely novel target in cancer therapeutics.

**Figure 1:**
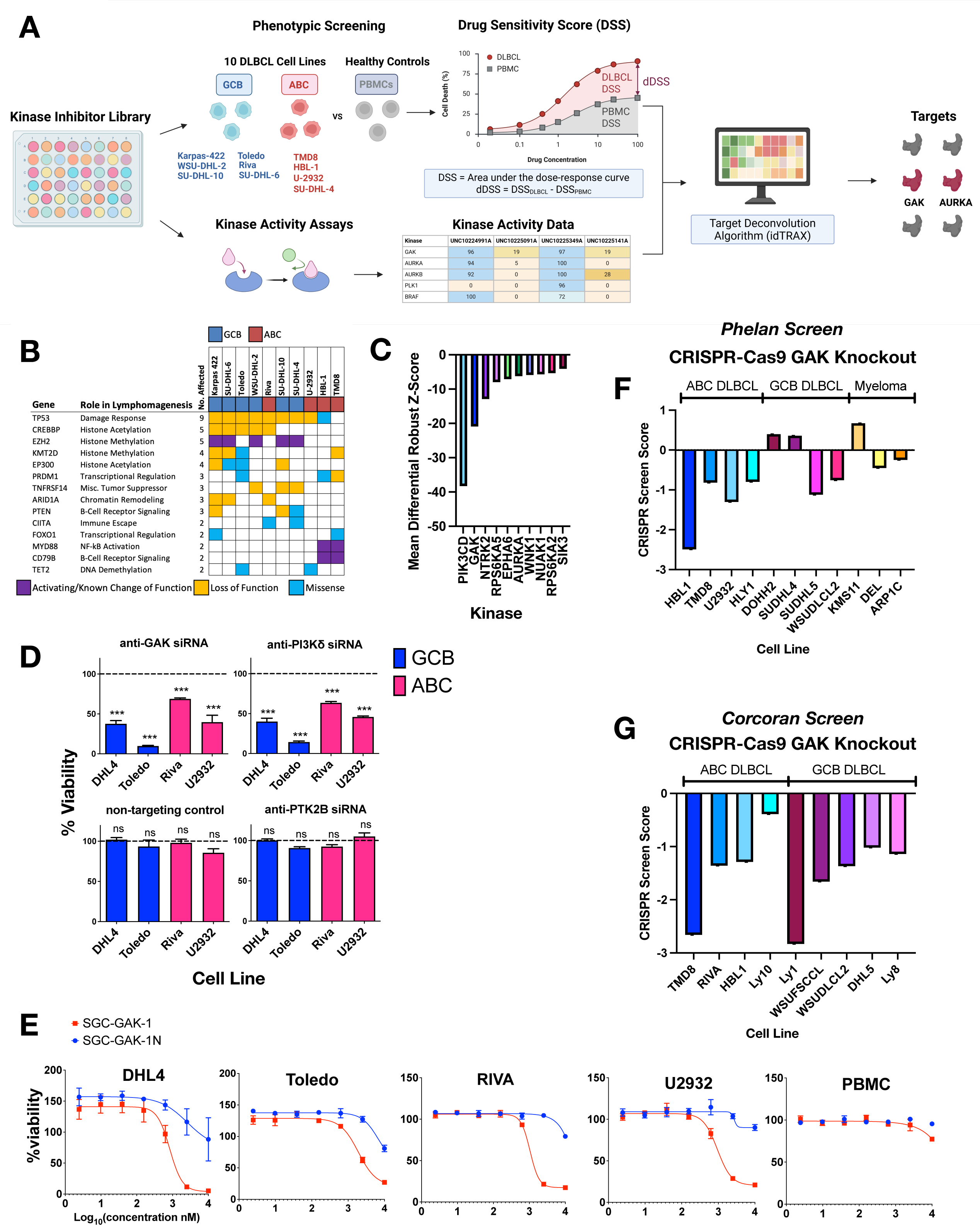
Phenotypic screen reveals GAK as a novel target for DLBCL. **1A:** Schematic of the multistep target discovery pipeline used to identify novel therapeutic targets in DLBCL. **1B:** Altered genes in 10 DLBCL lines in the screen, selected to capture genomic diversity. **1C:** Mean differential robust z-scores of siRNAs to validate hits from screen. The differential score reflects relative degree of tumor selectivity for each target. **1D:** Percent viability of DHL4, Toledo, RIVA, and U2932 cells following siRNA-mediated depletion of GAK, PI3Kδ, PTK2B, and a non-targeting control compared to parental cell lines. Data represent mean ± SEM from four technical replicates. ****p<0.0001 compared to untreated. **1E:** Dose response viability assays of DHL4, Toledo, RIVA, and U2932 cells treated with SGC-GAK-1 and SGC-GAK-1N for 72 hours. Data represent mean ± SEM from four technical replicates. **1F-G:** GAK dependency across ABC and GCB subtype DLBCL and Myeloma cell lines in independent CRISPR-Cas9 screens. CRISPR screen scores (CSS) represent the standardized effect of gene knockout. Strongly negative CSS values indicate GAK is a lethal dependency in all models (Phelan, F; Corcoran, G).

To validate idTRAX results, we assessed siRNA knockdown of the top 10% of hits in two GCB and two ABC cell lines, confirming AURKA and GAK **(Fig. 1C, Table S6-7)**. We then performed a second siRNA confirmation limited to GAK, PI3Kδ (positive control), PTK2B (negative screen hit), and a non-targeting control (siNT, **Fig. 1D)**. Si*GAK* consistently promoted decreased viability in DLBCL cells. Because GAK has not previously been highlighted as a potential target in lymphoma, we examined data from published genome-wide CRISPR-Cas9 dependency datasets. Across two independent CRISPR knockout screens, GAK loss produced strongly negative dependency scores in both ABC and GCB DLBCL models, consistent with a lethal dependency **(Fig. 1F-G)**.^22,23^ Therefore, both *GAK* RNAi depletion and genetic ablation are sufficient to drive DLBCL cell death.

Since these initial validation steps did not assess importance of GAK’s kinase function, we next employed the selective ATP-competitive GAK inhibitor SGC-GAK-1, alongside its GAK-sparing analog SGC-GAK-1N.^24^ SGC-GAK-1 demonstrated concentration-dependent killing across DLBCL cell lines but spared PBMCs, while the GAK-sparing analog showed minimal potency **(Fig. 1E)**. SGC-GAK-1 has reported off-target affinity for RIPK2, a liability not shared by SGC-GAK-1N.^24^ However, interrogation of CRISPR dependency datasets demonstrated RIPK2 ablation does not impair DLBCL cell viability, in contrast to the strong dependency observed for GAK **(Fig. S1B-C)**.^22,23^ Therefore, these data indicate that the observed cytotoxicity arises from inhibition of GAK rather than off-target RIPK2 suppression.

Collectively, a combination of phenotypic screening and target deconvolution, followed by validation with RNAi, CRISPR screening results, and chemical inhibition revealed GAK as a novel target candidate in DLBCL.

### GAK Inhibition is Broadly Active Against DLBCL Without Consistent Impact on B-Cell Receptor Signaling or Internalization

We undertook broader assessment with SGC-GAK-1 (GAKi) against a large panel of DLBCL lines **(Fig. 2A)**. In growth inhibition assays, the compound showed ≤1 µM GI_50_ across systems. Consistent with CRISPR results, ABC-derived lines were generally more sensitive than those derived from GCB cases (p=0.0078) **(Fig. 2B)**. Various other hematologic and solid-tumor malignancies also demonstrated sensitivity **(Fig. S2A)**. GAK is a clathrin heavy chain (CHC) binding partner and component of clathrin-coated vesicles during endocytosis, a process whose disruption may alter oncogenic signals.^13,15,25^ B-cell receptor (BCR) signaling drives DLBCL pathogenesis and employs trafficking to recycle BCR components to the cell surface after activation-induced endocytosis and to form MYD88-TLR9- BCR (My-T-BCR) supercomplexes in endolysosomes.^26,27^ Given GAK’s role in vesicular trafficking, we investigated whether its inhibition disrupts BCR signaling or internalization. We stimulated DLBCL cells with soluble α-IgM to activate BCR signaling through receptor crosslinking. As expected, downstream pathways were robustly activated, evidenced by increased phosphorylation of BTK, AKT, and ERK **(Fig. 2C).** Pretreatment with SGC-GAK-1 did not substantially alter these activation profiles, causing only minor attenuation of phospho-signals, which was inconsistent across systems. Instead, both stimulated and unstimulated conditions in all lines caused pronounced induction of phosphorylated histone 3 (pH3) in response to treatment, consistent with G2/M arrest (investigated further below). We next examined BCR internalization following receptor activation. DLBCL cells were pretreated with drug or DMSO for 2 hours, stimulated with α-IgM to induce receptor crosslinking, and subjected to a time-course analysis of BCR internalization. As expected, DMSO-treated control cells demonstrated a progressive reduction in surface BCR over time, while dynasore, a dynamin inhibitor blocking clathrin-mediated endocytosis (CME), inhibited receptor internalization, increasing BCR cell-surface retention **(Fig. 2D, Fig. S2B)**. In the Ly1 model, dynasore did not fully abrogate receptor internalization **(Fig. S2C-D)**, suggesting relative resistance to dynamin inhibition and consistent with prior observations that BCR endocytosis may also proceed through dynamin-independent endocytic pathways.^28^ Cells treated with SGC-GAK-1 closely mirrored DMSO controls, exhibiting comparable levels of BCR surface loss following stimulation **(Fig. 2D)**. Pharmacologic inhibition of GAK therefore did not impact BCR internalization in response to crosslinking.

**Figure 2:**
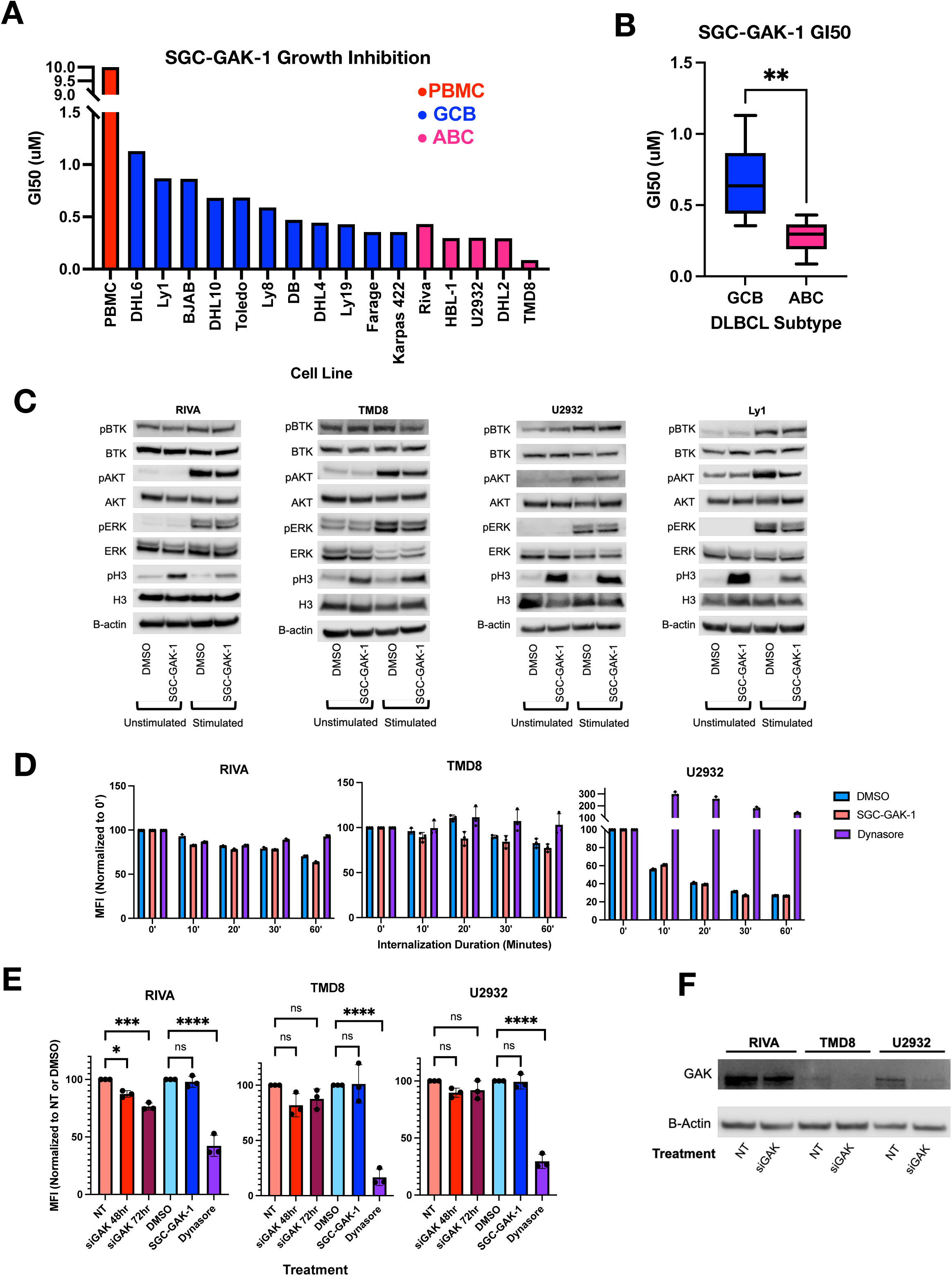
GAK inhibition is broadly suppresses growth in DLBCL and exerts anti-lymphoma effects independent of BCR signaling and trafficking. **2A:** GI_50_ values calculated from growth inhibition assays of a panel of GCB and ABC DLBCL cell lines and PBMCs treated with SGC-GAK-1. **2B:** Box plot of GI_50_ values comparing GCB and ABC DLBCL cell lines treated with SGC-GAK-1. p = 0.0078 **. **2C:** Immunoblot analysis of BCR signaling components in DLBCL cells treated with SGC-GAK-1 (1 µM, 6 hr) with or without α-IgM stimulation (10 µg/mL, 20 min). **2D:** Time course of BCR internalization measured by flow cytometric quantification of surface BCR (MFI) following α-IgM stimulation in DLBCL cells pretreated with DMSO, SGC-GAK-1 (1 μM), or dynasore (100 μM) for 2 hours. Surface IgM was detected using FITC-conjugated secondary antibody, and values were normalized to the 0-minute time point. **2E:** Transferrin uptake measured by flow cytometric MFI of CF®488A-conjugated transferrin in DLBCL cells following pretreatment with DMSO, SGC-GAK-1 (1 μM), or dynasore (100 μM), or transfection with siGAK or non-targeting (NT) siRNA. Cells were serum-starved, incubated with labeled transferrin at 4°C for surface binding, and allowed to internalize upon warming to 37°C. MFI values were normalized to DMSO or NT controls as indicated. **2F:** Immunoblot analysis of GAK protein levels in DLBCL cells following transfection with siGAK or NT siRNA.

GAK’s roles in endocytosis and vesicular trafficking have been defined primarily through RNAi knockdown loss-of-function approaches, while the role of its kinase function is minimally characterized. Notably, recent studies suggested GAK’s kinase activities may be dispensable to its participation in CME.^29^ To explain our results, we hypothesized GAK kinase activity is dispensable for endocytosis and instead essential for a different process whose inhibition resulted in G2/M arrest in DLBCL cells (**Fig. 2C**). To directly compare the effects of kinase inhibition versus protein depletion, we quantified transferrin uptake, a well-established readout of CME. Cells were pretreated with SGC-GAK-1, dynasore, or DMSO, or transfected with siGAK or non-targeting (NT) siRNA, followed by incubation with fluorescently labeled transferrin (Tf CF®488) to assess internalization. Transferrin was allowed to internalize upon warming, and uptake was quantified by flow cytometry as mean fluorescence intensity (MFI), normalized to DMSO or NT controls. As expected, dynasore treatment reduced transferrin uptake, confirming effective inhibition of CME **(Fig. 2E)**. In contrast, SGC-GAK-1 treated cells maintained transferrin uptake comparable to DMSO controls **(Fig. 2E)**. Consistent with BCR internalization results, the Ly1 cell line exhibited relative resistance to dynamin inhibition, but again there was no difference between DMSO and GAKi **(Fig. S2F)**. We next evaluated the effect of siRNA-mediated GAK depletion. In RIVA cells, GAK knockdown resulted in a reduction of transferrin uptake, whereas minimal impact was observed in additional models **(Fig. 2E-F, Fig. S2E).**

Despite variability in the effects of GAK depletion, pharmacologic GAK inhibition did not impair transferrin internalization in any model tested. Together, these results indicate that the broad anti-DLBCL activity of GAK inhibition is unlikely to arise from disruption of CME.

### GAK Inhibition Activates Spindle-Assembly Checkpoint Gene-Expression Responses

To assess the cellular impact of GAK kinase inhibition in a more unbiased fashion, we interrogated gene expression changes through RNA sequencing (RNA-seq, **Fig. 3A**). We isolated RNA from U2932 (ABC) and OCI-Ly1 (GCB) cells treated with either DMSO or 1 µM SGC-GAK-1 for 12 h or 24 h. Unsupervised clustering showed clear separation between SGC-GAK-1 and DMSO **(Fig. S3A-B)**. In Ly1, 100 and 749 genes were differentially expressed (p < 0.05, log-2 fold change > 0.5) at 12 and 24 hours respectively compared to DMSO (**Fig. 3B**). Similarly, 85 and 412 genes were differentially expressed in U2932 at 12 h and 24 h, respectively (**Fig. 3C**). Gene-set enrichment analysis by FGSEA^30^ revealed striking upregulation of sets associated with cell-cycle progression following GAK inhibition at both timepoints in both lines **(Fig. 3D-E, Fig. S3C-D)**. Closer examination revealed upregulation at both timepoints of multiple genes encoding spindle assembly checkpoint (SAC) participants, including *BUB1, BUB1B, CCNB1, AURKB*, and *PLK1* (**Fig. 3F-G, Fig. S3E-F)**, consistent with the induction of pH3 that we observed (**Fig. 2C**). Previous studies employing RNAi depletion implicated GAK in centrosome integrity and alignment of mitotic spindles during M phase but did not assess its kinase function in these activities.^14,16^ Our results therefore support a model in which GAK’s kinase activity regulates mitotic progression by DLBCL cells, and its inhibition is sufficient to trigger checkpoint activation and impair viability.

**Figure 3:**
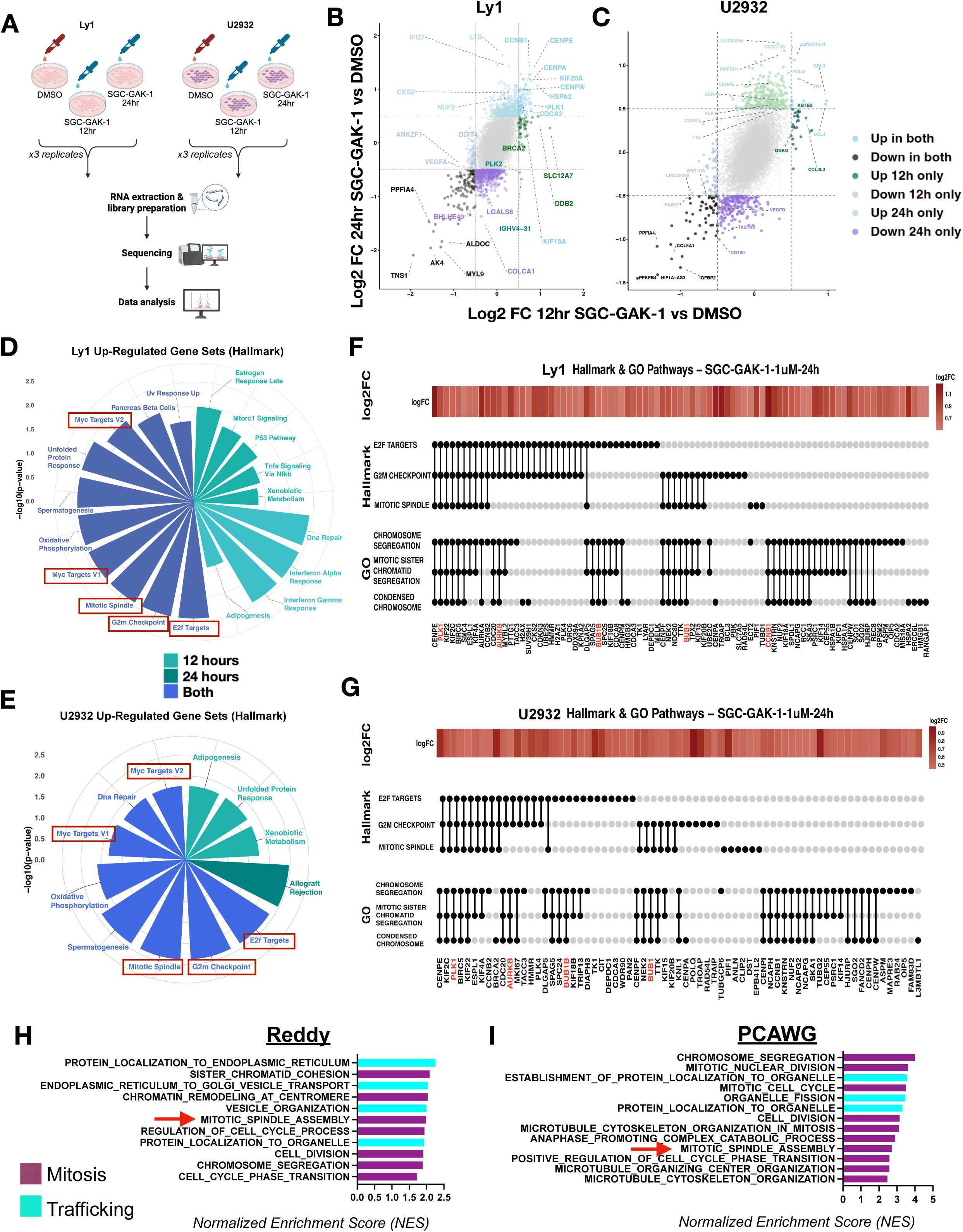
GAK Drives Mitotic Progression and Cell Cycle Activation in DLBCL. **3A:** Schematic of RNA sequencing workflow. U2932 (ABC) and Ly1 (GCB) DLBCL cells were treated with SGC-GAK-1 (1 μM) or DMSO for 12 and 24 hours in biological triplicate, followed by RNA isolation and transcriptomic analysis. **3B-C:** Differential gene expression following SGC-GAK-1 treatment (1 μM) for 12 and 24 hours vs DMSO. Genes meeting significance thresholds (p < 0.05, log2 fold change > 0.5) are colored by temporal regulation (both time points, 12 hours only, or 24 hours only). (B) Ly1; (C) U2932. **3D-E:** FGSEA of Hallmark pathways following SGC-GAK-1 treatment (1 μM) for 12 and 24 hours. Significantly enriched gene sets are shown and annotated by time point of enrichment (12 hours, 24 hours, or both). Gene sets associated with cell-cycle progression are highlighted. (D) Ly1; (E) U2932. **3F-G:** Genes within enriched cell cycle–associated gene sets following SGC-GAK-1 treatment (1 μM) at 24 hours. Spindle assembly checkpoint (SAC) genes are highlighted in red. Gene sets are derived from pathways identified in Fig. 3D–E. (F) Ly1; (G) U2932. **3H-I:** (H) Reddy and (I) PCAWG data show enriched genesets in *GAK*-high vs *GAK*-low cases from FGSEA analyses (adjusted p<0.05, log-2 fold change >0.5).

To further assess GAK in disease biology, we analyzed DLBCL gene-expression profiling (GEP) datasets from clinical cases. We divided cases from the Reddy RNA-seq dataset (n=775) into four quartiles based on *GAK* expression.^31^ Comparing *GAK*-high (Q4) to *GAK*-low (Q1) cases, we found 1,501 differentially expressed genes at adjusted p<0.05, log-2 fold change >0.5. Gene-set enrichment analysis by FGSEA^30^ revealed 34 gene sets upregulated in Q4 vs. Q1 at strict false discovery rate (q<0.05). Strikingly, 11 of these were directly relevant to either mitosis or vesicular trafficking **(Fig. 3H)**. For validation, we examined 75 DLBCL cases in the Pan-Cancer Analysis of Whole Genomes (PCAWG) data with available RNA-seq.^32^ Despite the relatively lower power of the smaller dataset, there were 599 differentially expressed genes between *GAK* Q4 and Q1. FGSEA strongly validated convergence on mitosis and trafficking functions in 13/60 gene sets at q<0.05 **(Fig. 3I)**. Both datasets implicate MITOTIC_SPINDLE_ASSEMBLY as strongly activated in *GAK*-high cases (red arrows). Based on the observed perturbation of mitotic gene expression in response to SGC-GAK-1’s block at G2/M, we investigated potential for synergistic interactions between SGC-GAK-1 and other drugs with antimitotic activities. We tested SGC-GAK-1 combined with DLBCL chemotherapeutics and various targeted cell-cycle inhibitors. We assessed 72-hour cell viability effects by SynergyFinder Bliss Synergy Score, indicating synergy at >10, additivity ≤10 to -10, or antagonism <-10. Strikingly, the combination treatments showed frequent antagonism across combination approaches and cell lines, and synergy was absent in all **(Fig. S3G)**, suggesting GAK inhibition alone profoundly disrupts essential cell cycle processes, rendering additional inhibitors redundant or counterproductive. Overall, these data implicate GAK kinase function as a key regulator of mitotic progression by DLBCL tumors.

### GAK Inhibition Triggers Induction of the Mitotic Spindle Assembly Checkpoint and G2/M-Phase Cell-Cycle Arrest

To further assess mitotic impact of GAK inhibition, we performed immunoblotting for key cell cycle regulators identified by RNA-seq. Treatment with SGC-GAK-1 led to increased expression of BUB1B, BUB1, and Cyclin B1 in parallel with pH3 induction **(Fig. 4A, Fig. S4A).**^33^ Notably, these effects were similar to those of vincristine, a vinca alkaloid chemotherapeutic that disrupts microtubule dynamics and promotes G2/M arrest **(Fig. 4A)**. Cell cycle assessment by quantitative DNA flow cytometry further revealed accumulation of SGC-GAK-1–treated cells in the G2/M phase at 24 hours, consistent with the arrest profile of vincristine **(Fig. 4B, Fig. S4B)**. These results further implicate GAK kinase activity in mitosis, aligning with its previously described, though poorly characterized, role in regulating mitotic spindles.^14,16^ Further assessment by immunofluorescent confocal imaging revealed pronounced distortion of spindles and misalignment of chromosomes in response to SGC-GAK-1 compared to vehicle **(Fig. 4C)**. Quantification of DAPI-stained chromosomes confirmed a significant increase in G2/M-phase cells, consistent with flow cytometry results and supporting mitotic arrest as a key phenotype. Notably, these effects were selective for malignant cells, as PBMCs were unaffected **(Fig. 4D)**. We also performed γ-tubulin immunofluorescence and observed multi-aster (more than two centrosomes) formation in SGC-GAK-1–treated DLBCL cells, indicated by multiple discrete γ-tubulin foci instead of a focused bipolar spindle **(Fig. 4E-F).**

**Figure 4:**
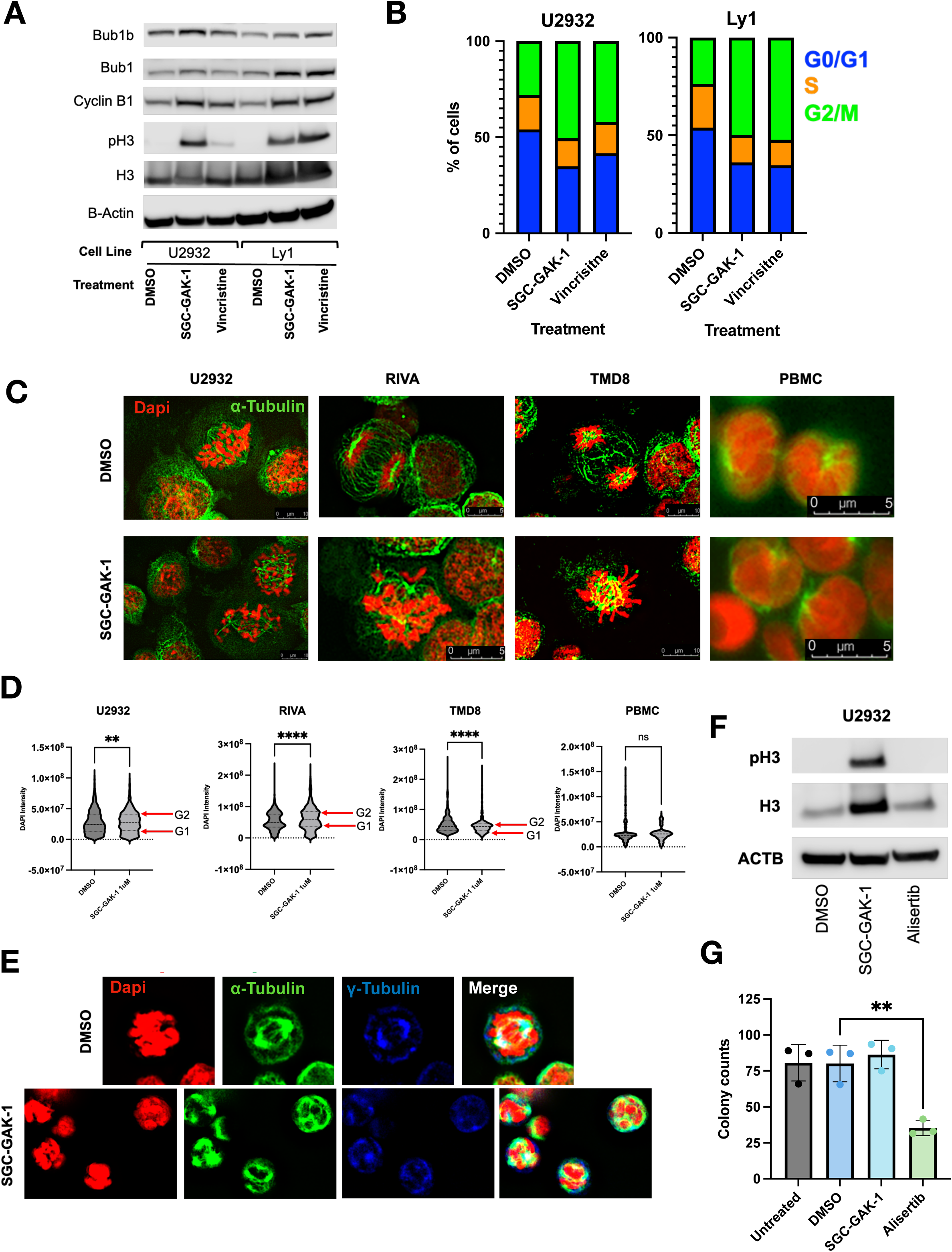
GAK inhibition triggers induction of the mitotic spindle assembly checkpoint and G2/M-phase cell-cycle arrest. **4A:** Immunoblot analysis of spindle assembly checkpoint and cell-cycle regulators in U2932 and Ly1 cells treated with SGC-GAK-1 (1 µM), Vincristine (1 nM), or DMSO control for 24 hrs. **4B:** Cell cycle distribution determined by flow cytometry (DNA content analysis) in U2932 and Ly1 cells treated with SGC-GAK-1 (1 μM), vincristine (1 nM), or DMSO for 24 hours. **4C:** Confocal immunofluorescence imaging of U2932, RIVA, TMD8, and PBMCs treated with SGC-GAK-1 (1 µM) or DMSO for 24 hrs. Chromosomes stained red; α-tubulin stained green. **4D:** Violin plots depicting quantification of DAPI-stained chromosomes following 24-hour treatment with SGC-GAK-1 (1 µM) versus DMSO control. ****p < 0.0001, **p < 0.01. **4E:** Representative confocal immunofluorescence images of RIVA cells treated with SGC-GAK-1 (1 µM) or DMSO for 24 hrs, highlighting multi-aster formation with γ-tubulin staining. **4F:** Immunoblot analysis of phospho-histone H3 (pH3) and total H3 in U2932 cells treated with SGC-GAK-1 (1 μM), alisertib (1 μM), or DMSO for 24 hrs. **4G:** Colony counts from a colony-forming unit (CFU) assay of bone marrow cells from wild-type C57BL/6 mice following treatment with SGC-GAK-1 (1 μM), alisertib (1 μM), DMSO, or untreated control.

This abnormal spindle architecture is similar to previously reported effects of GAK depletion.^16^ These data implicate GAK kinase function as a key participant in mitotic progression, leading to spindle atypia, G2/M arrest, and cell death upon targeted inhibition.

We next compared GAK inhibition to AURKA inhibition with alisertib, a drug that failed to progress as a DLBCL therapeutic due to high rates of hematologic toxicities.^21^ AURKA has well-defined roles at multiple cell-cycle phases, and previous gene-expression analyses assessing alisertib showed down-regulation of cell-cycle factors at the level of mRNA expression as part of an overall cellular response,^34^ in marked contrast to our SGK-GAK-1 results (**Fig. 3D-E**). We hypothesized GAK inhibition’s impact is more focal, further supported by western blotting, showing pH3 induction by SGC-GAK-1 that was not induced by an equipotent dose of alisertib (**Fig. 4F**). Because GAK emerged from a screening pipeline specifically designed to reveal targets with tumor-selective activities, we wanted to directly compare GAK and AURKA inhibition in a relevant non-malignant context. We therefore assessed SGC-GAK-1 and alisertib in colony-forming unit (CFU) assays on bone marrow cells from wild-type C57BL/6 mice.

While 1 µM alisertib eliminated greater that half of CFU capacity by marrow progenitors compared to DMSO (p=0.0025), 1 µM SGC-GAK-1 had no impact. We note 1 µM SGC-GAK-1 is well within efficacy range against DLBCL cell lines (**Fig. 4G**). GAK is therefore a mitotic kinase with distinct cellular response and relative sparing of hematopoietic stem cells compared to the well studied and so-far clinically unsuccessful target AURKA.

### Retinoblastoma Deficiency Carries Increased Sensitivity to Mitotic Disruption by GAK Inhibition

We next examined GAK within a broader context of DLBCL pathogenesis, in which cell-cycle activation is well-described, converging on inactivation of the tumor-suppressive master cell-cycle regulator retinoblastoma associated protein (RB) encoded by *RB1*.^35,36^ Though previous work in lung and breast cancers revealed RB-deficient tumors are more sensitive to AURKA or AURKB inhibition,^37,38^ subsequent CRISPR screening showed GAK loss-of-function is a synthetic lethal vulnerability in systems with *RB1* deletion.^39^ Neither AURKA nor AURKB were hits by this approach. We therefore tested for a relationship between *GAK* and *RB1* in DLBCL gene expression data, focusing chip-based data from Lenz et al. (n=414) and RNA-seq results in Reddy et al. (n=775).^3,31^ We divided the Lenz cohort based on *GAK* expression into GAK-high (n=182) and GAK-low (n=232). We compared quartiles for the quantitative Reddy RNA-seq data. Analysis of both datasets revealed strong association between low *RB1* expression and increased *GAK* expression **(Fig. 5A-B**; p=2.13e-22 and 2.2e-16 respectively). Examining cell lines, we found RIVA and U2932 cells completely lack RB **(Fig. 5C)**, and we note these lines had by far the most striking arrest at G2/M and mitotic atypia in response to SGC-GAK-1 **(Fig. 4B-D)**. We therefore further assessed these cells compared to RB-competent SU-DHL-4 cells using SGC-GAK-1 and SGC-GAK-1N, both at 10 µM. Flow cytometry showed pronounced G2/M arrest in RIVA and U2932 compared to DHL4 **(Fig. 5D, Fig. S5A)**, while the GAK-sparing 1N analog had no impact, again highlighting specificity of GAK kinase inhibition as driving these effects.

**Figure 5:**
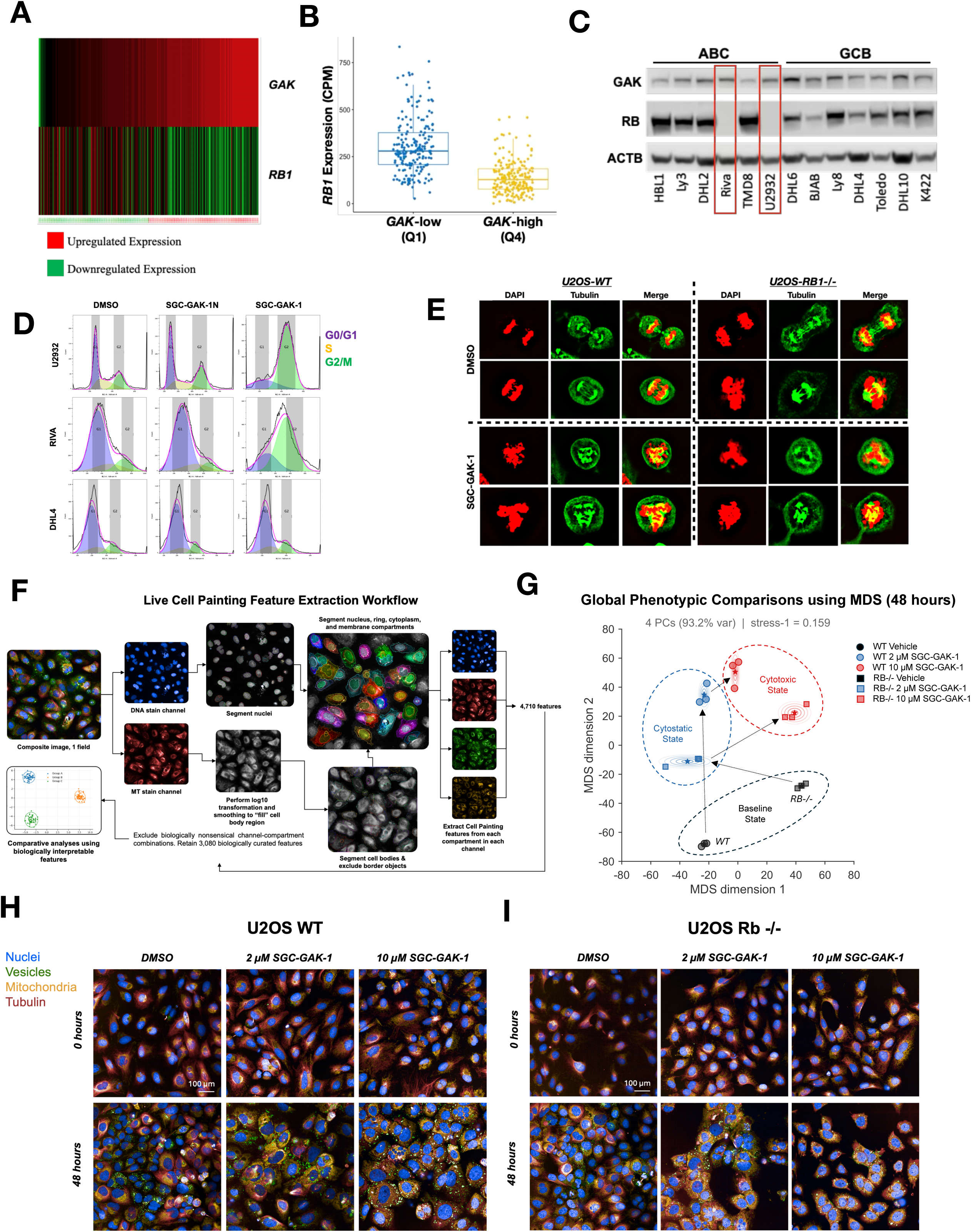
GAK dependence is strongly linked to retinoblastoma loss of function in DLBCL. **5A-B:** (A) Lenz and (B) Reddy data show low *RB1* associated with increased *GAK*. p=2.13e-22 *RB1* expression difference by *GAK* group (Lenz); p=2.2e-16 (Wilcoxon) Q1 vs Q4 (Reddy). **5C:** Immunoblot analysis of GAK and RB in DLBCL cell lines. **5D:** Flow cytometry histograms of DNA content (Vybrant DyeCycle Orange) used for cell cycle analysis in DLBCL cells treated with SGC-GAK-1 (10 μM), SGC-GAK-1N (10 μM), or DMSO for 24 hours. **5E:** Confocal immunofluorescence imaging of WT and RB1–/– U2OS cells treated with DMSO or SGC-GAK-1 (1 µM, 24 hr). Chromosomes stained red (DAPI), α-tubulin stained green. **5F:** Live Cell Painting workflow schematic. High-content live-cell imaging (Phenovue) was used to extract morphological features, which were analyzed by multidimensional scaling to define phenotypic states. **5G:** Multidimensional scaling of principal components derived from 3,080 Cell Painting features in U2OS WT and RB⁻/⁻ cells under indicated treatment conditions. **5H-I:** Representative Live Cell Painting images of U2OS WT (H) and RB⁻/⁻ (I) cells treated with SGC-GAK-1 (2 μM or 10 μM) or DMSO, at 0 and 48 hours.

Evidence implicating increased dependence on GAK in RB-deficient tumors requires validation in an independent system, preferably one suitable for high-resolution imaging studies, which are challenging in DLBCL cell lines that grow in suspension. We therefore employed U2OS human osteosarcoma cells commonly used in cell cycle studies, and we obtained a well-characterized CRISPR/Cas9-modified *RB1*-deficient clone (kind gift from F.A. Dick).^40^ Though GAK protein levels were similar between WT and RB-deficient U2OS, pH3 induction was enhanced in RB–/– at timepoints up to 24 h prior to the onset of cell death (**Fig. S5B**). Confocal imaging revealed that even in an absence of treatment, RB1-deficient cells displayed chromosomal abnormalities relative to WT **(Fig. 5E)**. Upon GAK inhibition, both cell types showed profound mitotic defects, including chromosome misalignment, disorganized spindles, multi-aster structures, and accumulation at G2/M as inferred from DAPI intensity quantifications **(Fig. 5E, Fig. S5C)**.

For time-course evaluation, we employed high-content live-cell imaging using Live Cell Painting (Phenovue), with features analyzed by multidimensional scaling to define phenotypic states **(Fig. 5F)**. Based on initial cell viability results **(Fig. S5D)**, WT and RB-deficient U2OS cells were treated with SGC-GAK-1 at 2 μM (∼IC_50_) and 10 μM (≥IC_90_) and imaged over 48 hours. Time-lapse nuclear analysis showed that 2 μM GAKi arrested proliferation in both genotypes, resulting in ∼50% fewer nuclei by 48 hours **(Fig. S5E)** while preserving per-cell ATP levels, consistent with cytostasis. In contrast, 10 μM treatment yielded similar nuclear counts but near-complete ATP loss, suggesting a cytotoxic effect. Principal component analysis revealed that control, 2 μM, and 10 μM conditions occupy distinct morphological spaces in both WT and Rb-deficient U2OS cells **(Fig. 5G-I)**. At baseline, RB-deficient cells were displaced from WT controls, consistent with underlying mitotic stress. Upon treatment, both genotypes shifted along a shared cytostatic trajectory; however, RB-deficient cells exhibited greater displacement toward cytotoxic states, consistent with enhanced sensitivity to GAKi. The complete timecourse of imaging for all six conditions are provided as .mp4 files (**Supplementary Movies S1-6**).

Feature-level analysis revealed pharmacodynamic and mechanistic effects **(Table S8-9**). GAKi induced dose-dependent redistribution of vesicle-associated signal, with reduced peripheral staining and increased intracellular accumulation, consistent with perturbation of vesicular trafficking. Nuclear and tubulin features revealed pronounced mitotic disruption: cells adopted a rounded morphology with dose-dependent increases in nuclear area, indicative of 4N DNA accumulation, and exhibited micronuclei formation from chromosome missegregation.

Tubulin organization was markedly disrupted, with loss of spindle symmetry and dose-dependent accumulation of signal into discrete cytoplasmic foci, consistent with spindle collapse and failed mitotic exit. Loss of cortical tubulin symmetry was also observed, consistent with disruption of astral microtubule positioning.^16^ Mitochondrial features showed dose-dependent fragmentation and puncta enlargement, consistent with metabolic stress secondary to prolonged mitotic arrest.

RB-deficient cells exhibited amplified responses to GAKi across multiple readouts at the cytotoxic dose of 10 μM **(Table S9)**.Tubulin aggregation and nuclear abnormalities, including chromatin disorganization, nuclear swelling, and micronuclei accumulation, were more pronounced in RB-/- cells, consistent with enhanced mitotic catastrophe. Mitochondrial disruption was similarly exacerbated. The baseline mitochondrial alterations in RB-deficient systems likely further sensitize these cells to metabolic stress induced by prolonged mitotic arrest.^41^ This heightened vulnerability is reflected in the steeper dose-response observed in RB-/-cells **(Fig. S5D)**, indicating a more abrupt transition from viability to cell death. Together, these results reveal the extensive impact of GAKi on mitosis and cellular integrity and highlight the differential sensitivity of RB-deficient cells.

### Multiple Clinical Kinase Inhibitors are Potent Against GAK, Underscoring Clinical Tractability and Providing Routes to Rapid Translation

No therapeutic GAK inhibitor candidate suitable for further development have yet been developed. High GAK selectivity of SGC-GAK-1 makes it an ideal probe in vitro.^24^ However, this compound is severely limited in vivo due to rapid cytochrome P450 enzyme metabolism (mouse liver microsomes t_1/2_ 5.7 min), leading to first-pass hepatic metabolism and poor pharmacokinetic (PK) properties (t_1/2_ <1 hr, C_max_ <100 ng/mL).^42^ Pretreatment with the broad spectrum P450 inhibitor 1-aminobenzotriazole (ABT, 50 mg/kg, oral gavage) previously extended SGC-GAK-1’s t_1/2_ to ∼3 hr (C_max_ 650 ng/mL).^42^ We therefore tested for in vivo anti-lymphoma activity of SGC-GAK-1 (10 mg/kg, oral gavage), with each dose given 2 hrs post-ABT, to NSG mice bearing luciferase-engineered U2932 tumors introduced by tail-vein injection. Compared to vehicle-treated animals, SGC-GAK-1 achieved significant tumor burden reduction (p=0.0003) **(Fig. S6A-B)**. Immunohistochemistry (IHC) of tumors that arose in lung and liver showed marked decrease in the Ki67 proliferation marker in SGC-GAK-1 treated samples **(Fig. S6C)**. These results demonstrate in vivo efficacy for targeted GAK inhibition against DLBCL tumors since ABT treatment alone had no effect on DLBCL cell lines and also did not exhibit toxicity towards non-malignant PBMCs **(Fig. S6D)**.

Given SGC-GAK-1’s severe limitations as an in vivo probe, we sought clinically available kinase inhibitors suitable for repurposing against GAK. Review of published kinome profiling data revealed GAK is a off-target of multiple compounds, including several clinically-approved drugs.^43^ Strikingly, some of these have significantly higher binding affinity for GAK than their intended clinical targets **(Fig. 6A)**.^44^ Given extensive clinical use of these compounds, these data suggest GAK inhibition may be well tolerated in patients, and we obtained them for further investigation. NanoBRET live-cell target engagement assays showed that intracellular GAK potency by OTS167, milciclib, bosutinib, gilteritinib, and fedratinib was even higher than for SGC-GAK-1 **(Fig. 6B)**. Against DLBCL, Milciclib’s GI_50_ was 0.2-0.3 uM across GCB and ABC cell lines **(Fig. 6C)**. OTS167, whose GAK Kd_app_, as shown by nanoBRET, was over two orders of magnitude more potent than SGC-GAK-1 at 1.4 nM, demonstrating GI_50_ <10 nM across multiple lines **(Fig. 6D)**. Both OTS167 and milciclib promoted pH3 accumulation and G2/M cell cycle arrest **(Fig. 6E-G).** Stronger G2/M arrest at lower concentrations by OTS167 is consistent with its exquisitely high intracellular GAK potency, leading to the effects of hitting that target, compared to multi-target effects at higher concentrations. These findings provide orthogonal GAK target validation and highlight the potential for rapid clinical translation by repurposing existing kinase inhibitors.

**Figure 6:**
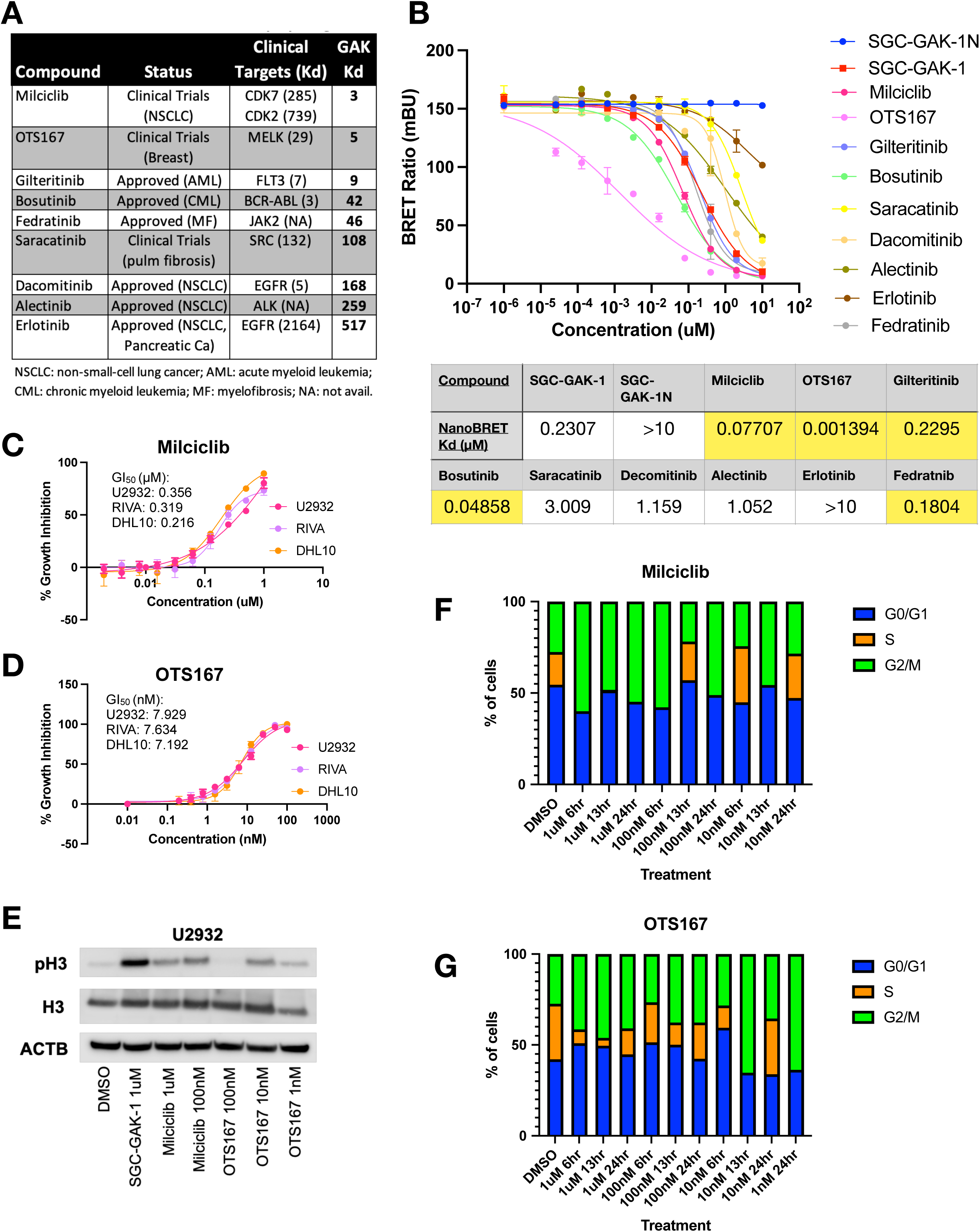
Candidate clinical drugs for GAK repurposing. **6A:** Biochemical affinities (K*_d_*)for GAK and intended clinical targets of clinically-approved compounds with reported off-target activity against GAK. **6B:** NanoBRET® target engagement (TE) intracellular kinase assay dose–response curves for SGC-GAK-1, SGC-GAK-1N, and clinically-approved compounds in transfected 293T cells. Calculated NanoBRET (K*_d_*) values are shown, including compounds with greater potency than SGC-GAK-1 (highlighted). **6C-D:** Dose–response viability assays of U2932, RIVA, and DHL10 cells treated with Milciclib (C) or OTS167 (D) for 72 hours. Data represent mean ± SEM of technical replicates. **6E:** Immunoblot analysis of cell cycle regulators, including histone H3 and phospho-histone H3 (pH3), in U2932 cells treated with SGC-GAK-1 (1 μM), Milciclib (1 μM, 100 nM), or OTS167 (100 nM, 10 nM, 1 nM) for 24 hours. **6F-G:** Cell cycle distribution determined by flow cytometry in U2932 cells treated with Milciclib (F) or OTS167 (G) at indicated concentrations (1 μM, 100 nM, 10 nM, 1 nM) for 6, 13, and 24 hours, compared to DMSO control.

### Clinical-stage OTS167 Demonstrates In Vivo Efficacy in a DLBCL PDX Model

OTS167 is an orally bioavailable, clinical-stage small molecule initially developed to target MELK in breast, lung, prostate, and pancreas cancers.^45^ However, kinome profiling data and our experimental results demonstrate that this compound inhibits GAK with greater selectivity than MELK (**Fig. 6A**).^43^ OTS167 has been tested in Phase I and Phase II clinical trials for solid tumors and leukemias, where it has shown favorable tolerability, supporting its potential for rapid repurposing.^46–48^ To assess its in vivo efficacy against DLBCL, we employed a PDX model derived from treatment-naïve GCB DLBCL (75549R2), with tumors implanted subcutaneously into NSG mice (**Fig. 7A**). Single-agent OTS167 led to a significant reduction in tumor volume (p < 0.0001), improved overall survival (p = 0.06), and a decrease in tumor weight evaluated ex vivo at endpoint (p = 0.09) compared to vehicle **(Fig. 7B-D)**. IHC of tumor samples showed an increase of pH3 staining, demonstrating cell cycle arrest in OTS167 treated mine, with a decrease in the Ki67 proliferation marker (**Fig. 7E**). These results confirm that a potent pharmacologic GAK inhibitor is effective in suppressing tumor growth in vivo as a single agent and support further investigation of OTS167 as a repurposed therapeutic in DLBCL.

**Figure 7:**
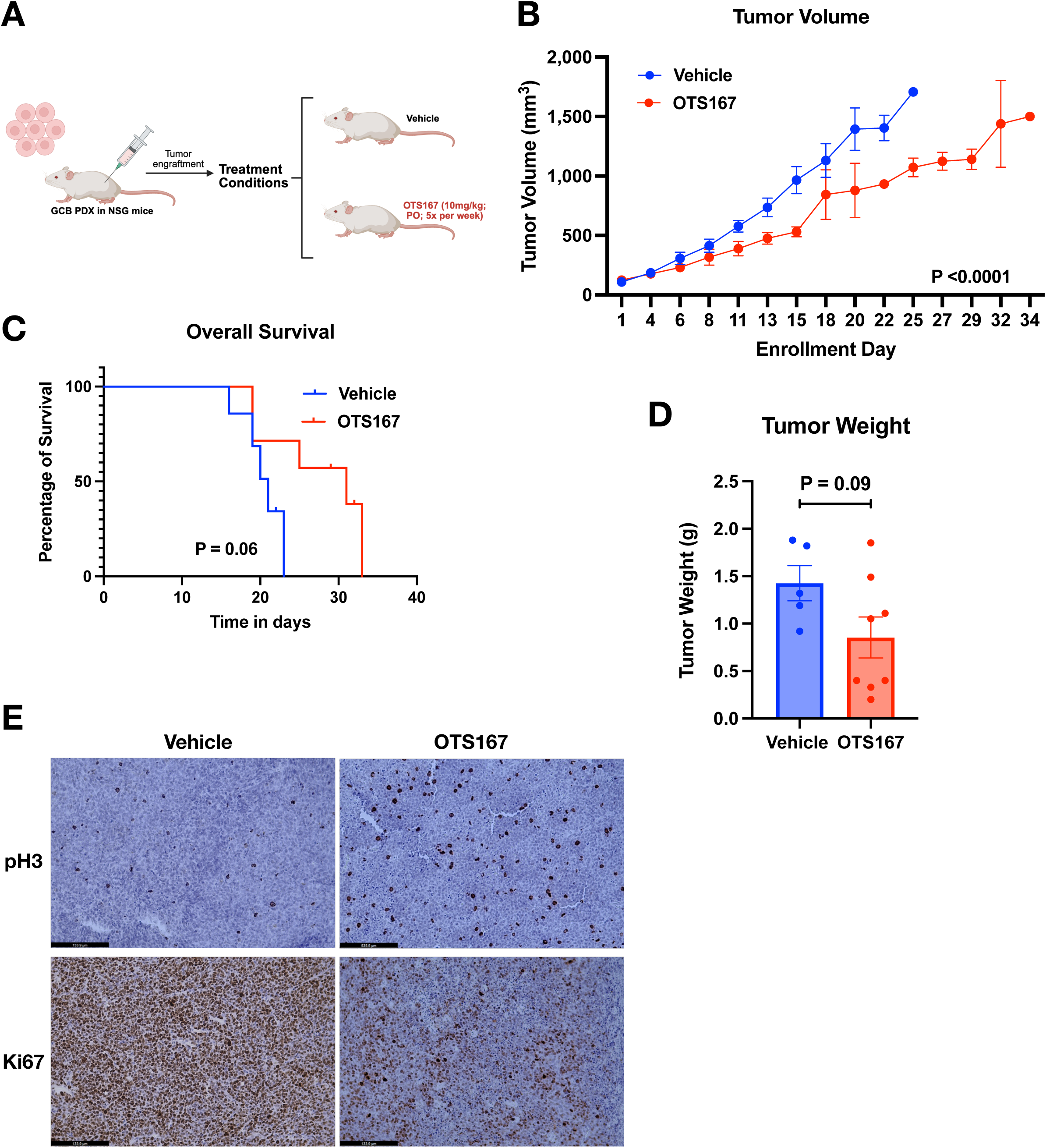
Clinical-stage OTS167 Demonstrates In Vivo Efficacy in a DLBCL PDX Model. **7A:** Schematic of experimental design for subcutaneous implantation and treatment of GCB-DLBCL PDX tumors (75549R2) in NSG mice with vehicle or OTS167 (10 mg/kg, oral gavage, 5×/week). **7B:** Tumor growth curves of NSG mice bearing GCB-DLBCL PDX tumors treated with vehicle or OTS167. Tumor volume was measured by ultrasound imaging; ***p < 0.0001 (two-way ANOVA). **7C:** Kaplan–Meier survival curves of mice from vehicle- and OTS167-treated groups (p = 0.06). **7D:** Tumor weights measured ex vivo at endpoint in vehicle- and OTS167-treated groups (p = 0.09, unpaired t test). **7E:** IHC staining for phospho-histone H3 (pH3) and Ki67 in tumor tissues from vehicle- and OTS167-treated mice.

## Discussion

Kinase-directed therapies have transformed the treatment of several B-cell non-Hodgkin lymphomas but have had limited impact in DLBCL. In chronic lymphocytic leukemia (CLL), BTK inhibition alone often provides durable disease control,^49^ largely replacing chemotherapy and marking a paradigm shift in clinical practice over the past decade and a half. In contrast, DLBCL’s molecular and clinical heterogeneity has defied such single-target approaches, and the frontline therapy with curative intent remains intensive combination chemoimmunotherapy: R-CHOP, pola-R-CHP, and variations thereof.^7^ Unfortunately, these approaches still fail in 30-40% of patients. ViPOR achieved impressive responses in rel/ref disease, including in patients with high-risk double-hit (*MYC* and *BCL2* rearranged) tumor cytogenetics, and without inclusion of any traditional chemotherapautic agents.^11^ The combination is not yet included in treatment guidelines, however, and single-agent ibrutinib is listed only as a palliative approach in non-transplant eligible rel/ref patients. Here we applied phenotypic screening and computational target deconvolution to reveal a novel kinase driver of DLBCL’s malignant biology, one available for rapid evaluation as a clinical target thanks to drug repurposing. Specifically, we find the cyclin-G associated kinase (GAK) is expressed in DLBCL cases associated with activation of mitotic gene-expression programs and with deficiency of *RB1*. Disruption of GAK kinase activity promotes G2/M mitotic arrest greater than any impact on GAK’s more well-described, though not clearly kinase-dependent, roles in endocytosis and vesicular trafficking. RB-deficient cells demonstrate the most rapid and pronounced onset of these effects, providing a readily available candidate biomarker for evaluation in clinical trials.

Previous studies showed that, like its binding partner CHC, GAK localizes to mitotic spindles, and that its siRNA knockdown promoted spindle positioning defects, multi-aster formation, and chromosomal atypia in post-mitotic cells.^13–16^ However, the potential role of its S/T kinase activities in cell cycle regulation remained unclear. We show that GAK kinase inhibition (GAKi) exhibits tumor-selective potency against DLBCL. Flow cytometry, western blotting, immunofluorescent confocal microscopy, and high-content live-cell imaging studies demonstrate mitotic disruption, leading to chromosome misalignment, spindle distortion, SAC activation, and cell cycle arrest. In contrast, GAK’s roles in endocytosis and vesicular trafficking in DLBCL appear limited. BCR internalization, for example, proceeded with comparable kinetics in the presence of GAKi, indicating that receptor trafficking following activation was largely preserved. Moreover, comparison between pharmacologic GAKi and siRNA-mediated depletion in a transferrin internalization assay demonstrated that CME was not consistently impaired across DLBCL models. While dynasore effectively blocked transferrin uptake, confirming assay sensitivity, SGC-GAK-1 did not reduce internalization relative to control conditions. Similarly, GAK depletion failed to significantly alter transferrin uptake in most cell lines tested, suggesting that the contribution of GAK to CME may vary across cellular contexts. We did observe vesicular compartment disruption by high-content live-cell imaging (cell painting), but these effects occurred in the context of multiple profound mitotic impacts that are more clearly associated with toxic cellular endpoints. Together, these findings indicate that disruption of CME is unlikely to account for the broad anti-DLBCL activity of GAK kinase inhibition. Instead, our results support a model in which the kinase-dependent functions of GAK contribute to mitotic integrity, and that pharmacologic inhibition of this activity induces mitotic catastrophe in lymphoma cells. The sparing of non-malignant cells further supports this conclusion, since these cells proliferate very slowly in culture while still carrying out endocytosis and trafficking as essential biologic processes. Encouragingly, we found GAKi did not impact hempatopoietic CFU formation in marked contrast to alisertib. Overall, our results establish for the first time that GAK is a cell cycle kinase whose inhibition results in anti-tumor activity.

This alone of course does not guarantee GAKi will be clinically tolerable; rapidly diving non-malignant cells, particularly of the hematologic system, could still be adversely impacted. Only dedicated clinical trials can truly establish therapeutic windows, but we provide compelling evidence that GAKi is likely tolerable in patients, specifically our observation that multiple drugs used as both approved cancer therapeutics as well as clinical trial cadidates exhibit potent off-target anti-GAK activity **(Fig. 6A)**. Notably, several of these bind to GAK more strongly than to their putative clinical targets, with higher GAK inhibitory activity than the selective tool compound SGC-GAK-1 **(Fig. 6B)**. These results are highly encouraging, suggesting that targeted GAK inhibition may be well-tolerated in patients. Moreover, repurposing these existing compounds can leverage established pharmacokinetics, dosing, and safety profiles. In particular, the clinical-stage molecule OTS167 offers a compelling opportunity. Although originally developed as a MELK inhibitor for solid tumors, kinome-profiling studies and our own experimental observations reveal OTS167 not only potently inhibits GAK, but also displays stronger selectivity toward GAK than MELK in biochemical and cellular assays. Its in vivo activity in our DLBCL PDX model provides preliminary evidence that a clinically advanced molecule can recapitulate the anti-lymphoma effects of GAK inhibition. Together, these features position OTS167 as an attractive proof-of-concept agent while medicinal chemistry efforts are underway to develop optimized GAK-directed therapeutics. Since single-agent non-immunotherapies have never found a role in curative-intent DLBCL clinical management, longer-term studies should focus rational combinations to most efficaciously leverage the impact of GAK inhibition.

Deeply defining the mechanisms by which GAK functions as a mitotic kinase, in particular the specific phosphorylation substrates that shape these activities, is beyond the scope of this study. However, we are currently analysing phosphoproteomics data of cells treated with and without GAKi in DLBCL and other cell systems. Preliminarily, these results paint a complex picture of candidate GAK substrates, each of which require extensive validation and functional characterization, studies that are outside our current scope. Overall, our research defines GAK as a novel and targetable dependency in DLBCL and opens new avenues for targeted therapy development. The frequent off-target inhibition of GAK by clinical kinase inhibitors including several approved drugs further supports feasibility and safety of inhibiting GAK as a therapeutic strategy.^43^ Future studies will continue to explore the molecular mechanisms underlying GAK’s role in DLBCL pathogenesis, in particular the specific phospho-substrates by which it regulates mitosis, and elucidating the broader impact of GAK inhibition on cancer biology. These efforts will deepen our understanding of DLBCL and contribute to the development of more effective, targeted therapies for this aggressive malignancy.

## Acknowledgements

This work was supported by grants from the NIH (1R41TR002293) to H.A., NIH/NCI (R01CA296318) to J.H.S. and H.A., Florida Department of Health (22B13) to J.H.S, H.A, Y.F., and S.S., and a Glaser Foundation Research Award (University of Miami) to H.A. Some experiments occurred at The Miami Project Drug Discovery Core (RRID:SCR_022542) and the Cancer Modeling and Onco-Genomic Shared Resources (CMSR RRID: SCR022891, OGSR RRID: SCR022502) of the Sylvester Comprehensive Cancer Center at the University of Miami, which is supported by the National Cancer Institute (NCI) of the

National Institutes of Health (NIH) under award number P30CA240139. The content is solely the responsibility of the authors and does not necessarily represent the official views of the NIH.

## Authorship Contributions

Contribution: O.B.F., B.A., P.M., A.K., P.K., L.L., A.D.N., M.L., D.H.P., P.A.Y., D.C., A.A., C.A.C., and H.A. performed bench experiments. S.V. and H.A. performed target deconvolution and other computational analyses. T.S., A.Y.A., O.B.F., and F.M. oversaw analysis of newly generated and published gene-expression data. M.G., B.S., and H.A. oversaw high-content image acquisition and data analysis. D.B. helped design and carry out in vivo experiments. A.B.M. performed in vivo imaging and efficacy experiments. S.C.S. and Y.F. helped design and analyze assessments of compound engagements with GAK. V.L. helped design and supervise phenotypic screening. J.H.S. and H.A. supervised all experiments. O.B.F., H.A., and J.H.S. wrote and edited the manuscript with input from all authors, and all authors have approved the final version of the manuscript. H.A. and J.H.S. supervised the project and are responsible for all content.

## Data Availability Statement

All data in this study were generated by the authors or were analyzed from primary sources cited and are available as needed upon request from the corresponding author. The RNA sequencing data we generated are publicly available from the Gene Expression Omnibus (https://www.ncbi.nlm.nih.gov/geo/, GEO) under accession# GSE279610.

