## Supplementary Figures for "The cyclin-G associated kinase (GAK) is a novel mitotic kinase and therapeutic target in diffuse large B-cell lymphoma"

### Supplementary Appendix

***SUPPLEMENTARY TABLES***

**Table S1.** Compounds used in cell-based experiments
**Table S2.** Antibody list
**Table S3.** Phenotypic screen results – DSS
**Table S4.** Phenotypic screen results – dDSS
**Table S5.** Target deconvolution results – idTRAX output
**Table S6.** siRNA validation – robust z-scores
**Table S7.** siRNA validation –differential robust z-scores
**Table S8.** Cell Paint feature heatmap
**Table S9.** Cell Paint detailed feature analysis

#### SUPPLEMENTARY TABLES 1-2 (all others in separate spreadsheet)

**Table S1.** Compounds/drugs in cell-based experiments

| S.no | Compound/drugs name | Source/catalog no. |
|---|---|---|
| **1** | SGC-GAK-1 | MedChemExpress, HY-122186 |
| **2** | SGC-GAK-1N | Sigma-Aldrich, SML2203 |
| **3** | Flavopiridol | MedChemExpress, HY-10005 |
| **4** | Alisertib | MedChemExpress, HY-10971 |
| **5** | Doxorubicin | MedChemExpress, HY-15142 |
| **6** | Gemcitabine | MedChemExpress, HY-17026 |
| **7** | Vincristine | MedChemExpress, HY-N0488 |
| **8** | Etoposide | MedChemExpress, HY-13629 |
| **9** | Cyclophosphamide | MedChemExpress, HY-17420 |
| **10** | Cisplatin | MedChemExpress, HY-17394 |
| **11** | Bendamustine | MedChemExpress, HY-B0077 |
| **12** | Dinaciclib | MedChemExpress, HY-10492 |
| **13** | Nocodazole | MedChemExpress, HY-13520 |
| **14** | Barasertib | MedChemExpress, HY-10127 |
| **15** | Volasertib | MedChemExpress, HY-12137 |
| **16** | Selinexor | MedChemExpress, HY-17536 |
| **17** | 1-aminobenzotriazole (ABT) | MedChemExpress, HY-103389 |
| **18** | Milciclib | MedChemExpress, HY-10424 |
| **19** | OTS167 | MedChemExpress, HY-15512 |
| **20** | Gilteritinib | MedChemExpress, HY-12432 |
| **21** | Bosutinib | MedChemExpress, HY-10158 |
| **22** | Fedratinib | MedChemExpress, HY-10409 |
| **23** | Saracatinib | MedChemExpress, HY-10234 |
| **24** | Dacomitinib | MedChemExpress, HY-13272 |
| **25** | Alectinib | MedChemExpress, HY-13011 |
| **26** | Erlotinib | MedChemExpress, HY-50896 |

**Table S2.** Antibody List

| S.No. | Antibody | Dilution | Application | Source/catalog no. |
|---|---|---|---|---|
| **1** | GAK | 1:100 | Western blot | CST, 51509 |
| **2** | AKT | 1:100 | Western blot | CST, 9272 |
| **3** | Phospho-AKT | 1:500 | Western blot | CST, 4060 |
| **4** | p44/42 MAPK (Erk1/2) | 1:1000 | Western blot | CST, 4695 |
| **5** | phospho-p44/42 MAPK (Erk1/2) | 1:500 | Western blot | CST, 4370 |
| **6** | GAPDH | 1:1000 | Western blot | CST, 5174 |
| **7** | Histone H3 | 1:1000 | Western blot | CST, 9715 |
| **8** | phospho-Histone H3 | 1:500 | Western blot | CST, 9701 |
| **9** | Cyclin B1 | 1:1000 | Western blot | CST, 4138 |
| **10** | BUB1B | 1:1000 | Western blot | CST, 4116 |
| **11** | β-Actin | 1:1000 | Western blot | CST, 3700 |
| **12** | RB | 1:1000 | Western blot | CST, 9313 |
| **13** | Anti-rabbit IgG, HRP-linked Antibody | 1:2000 | Western blot | CST, 7074 |
| **14** | Anti-mouse IgG, HRP-linked Antibody | 1:2000 | Western blot | CST, 7076 |
| **15** | Ki67 | 1:500 | IHC | CST, 9027 |

#### Figure S1

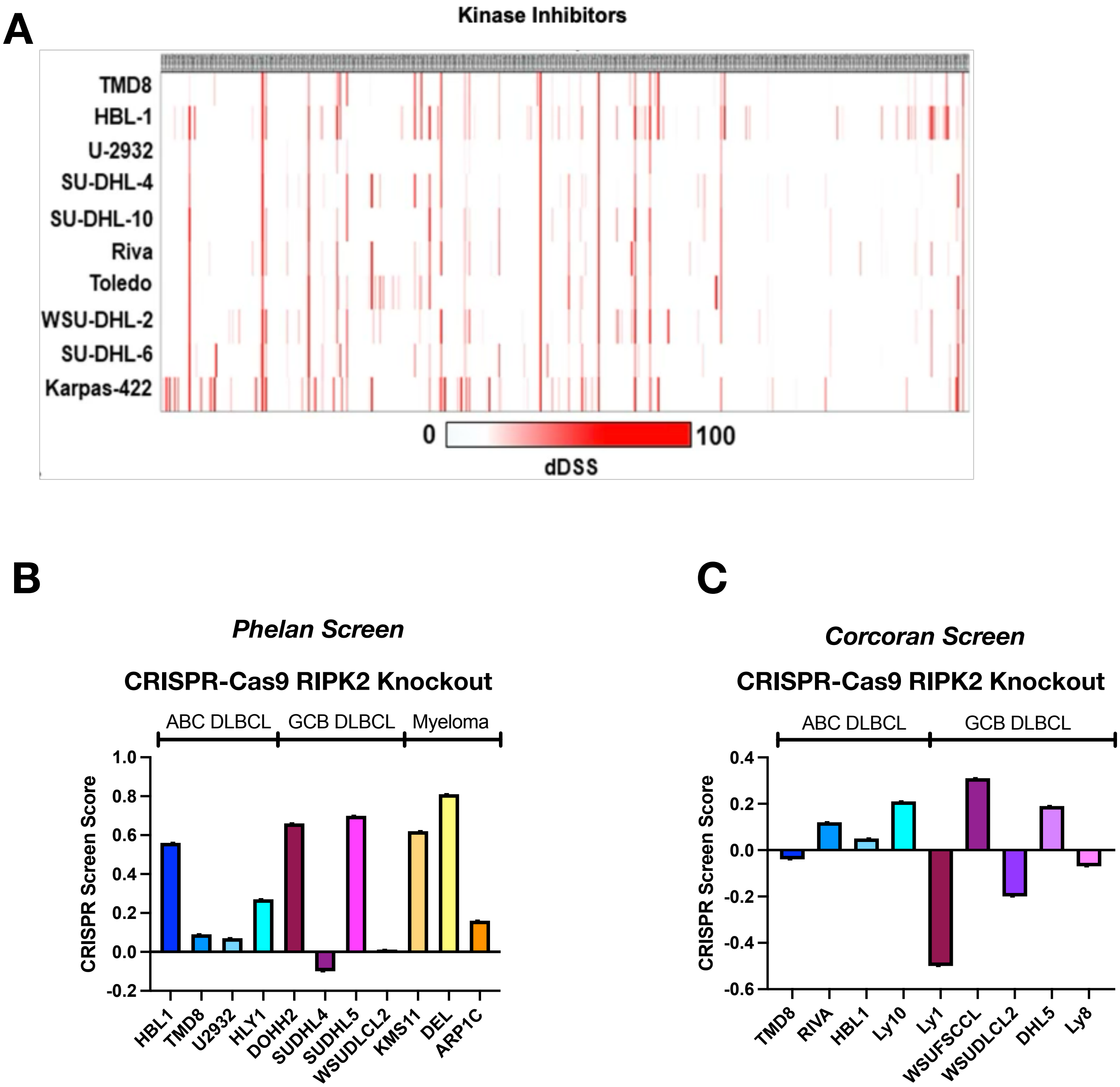

A
Kinase Inhibitors
TMD8
HBL-1
U-2932
SU-DHL-4
SU-DHL-10
Riva
Toledo
WSU-DHL-2
SU-DHL-6
Karpas-422
0
100
dDSS
B
Phelan Screen
CRISPR-Cas9 RIPK2 Knockout
ABC DLBCL
GCB DLBCL
Myeloma
CRISPR Screen Score
1.0
0.8
0.6
0.4
0.2
0.0
-0.2
HBL1
TMD8
U2932
HLY1
DOHH2
SUDHL4
SUDHL5
WSUDLCL2
KMS11
DEL
ARP1C
C
Corcoran Screen
CRISPR-Cas9 RIPK2 Knockout
ABC DLBCL
GCB DLBCL
CRISPR Screen Score
0.4
0.2
0.0
-0.2
-0.4
-0.6
TMD8
RIVA
HBL1
Ly10
Ly1
WSUFSCCL
WSUDLCL2
DHL5
Ly8

**Supplementary Figure S1** (previous page)

**S1A:** Differential drug sensitivity (dDSS) scoring of each kinase inhibitor against each cell line. Formula: dDSS = DSS (cell line) - DSS (PBMC).

**S1B-C:** RIPK2 dependency across ABC and GCB subtype DLBCL and Myeloma cell lines in independent CRISPR-Cas9 screens. CRISPR screen scores (CSS) represent the standardized effect of gene knockout (Phelan, B; Corcoran, C).

### Figure S2

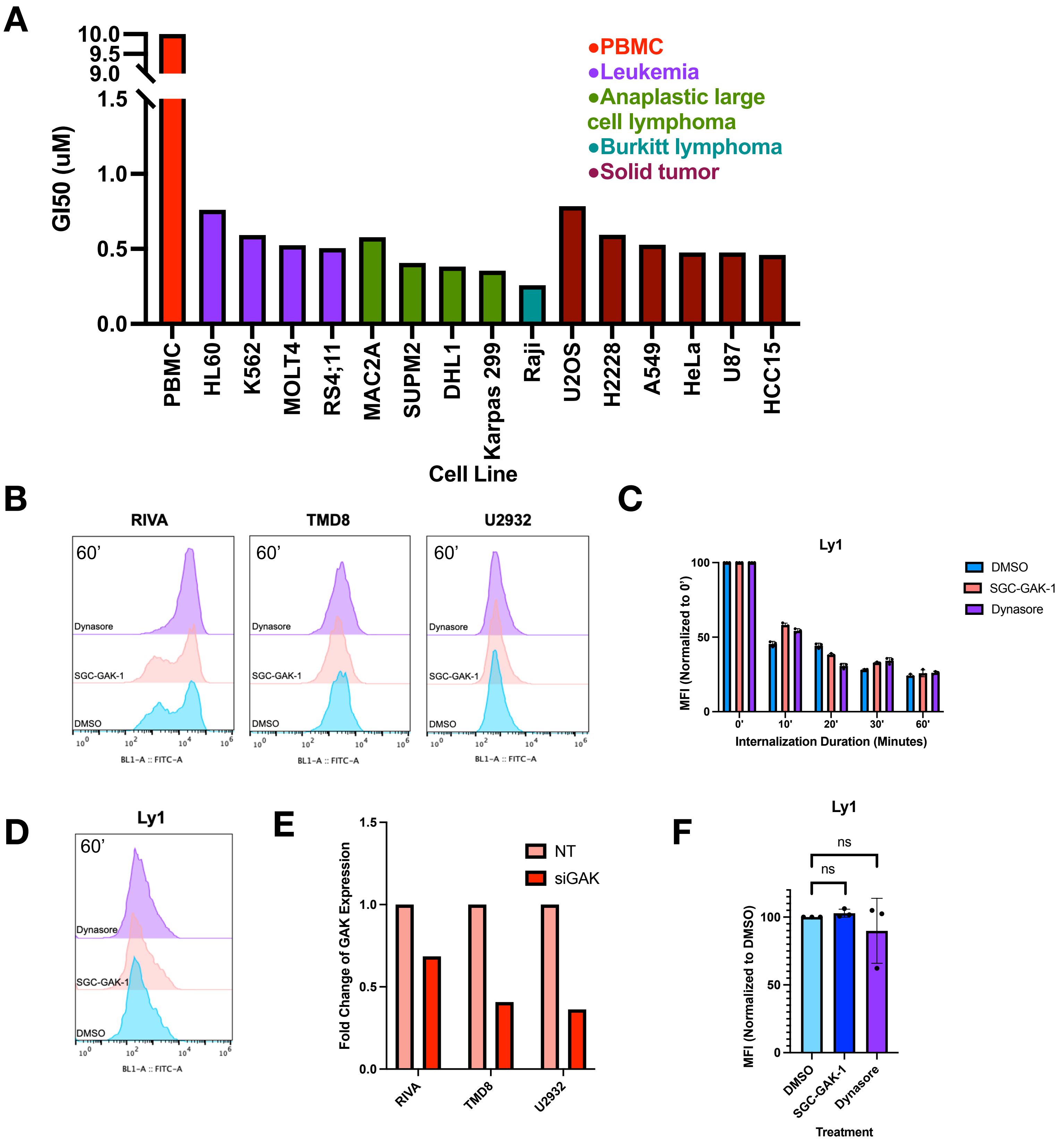
A
GI50 (uM)
10.0
9.5
9.0
1.5
1.0
0.5
0.0
●PBMC
●Leukemia
●Anaplastic large cell lymphoma
●Burkitt lymphoma
●Solid tumor
PBMC
HL60
K562
MOLT4
RS4;11
MAC2A
SUPM2
DHL1
Karpas 299
Raji
U2OS
H2228
A549
HeLa
U87
HCC15
Cell Line
B
RIVA
TMD8
U2932
60’
Dynasore
SGC-GAK-1
DMSO
BL1-A :: FITC-A
C
Ly1
MFI (Normalized to 0’)
DMSO
SGC-GAK-1
Dynasore
0'
10'
20'
30'
60'
Internalization Duration (Minutes)
D
Ly1
60’
Dynasore
SGC-GAK-1
DMSO
BL1-A :: FITC-A
E
Fold Change of GAK Expression
NT
siGAK
RIVA
TMD8
U2932
F
Ly1
ns
ns
MFI (Normalized to DMSO)
DMSO
SGC-GAK-1
Dynasore
Treatment

**Supplementary Figure S2** (previous page)

**S2A:** $GI_{50}$ values calculated from growth inhibition assays of leukemia, anaplastic large cell lymphoma, Burkitt lymphoma, and solid tumor cell lines treated with SGC-GAK-1.

**S2B:** Representative flow cytometry histograms of surface BCR (FITC signal) at the 60-minute time point following α-IgM stimulation in DLBCL cells pretreated with DMSO, SGC-GAK-1 (1 μM), or dynasore (100 μM), corresponding to Fig. 2D.

**S2C:** Time course of BCR internalization measured by flow cytometric quantification of surface BCR (MFI) following α-IgM stimulation in Ly1 cells pretreated with DMSO, SGC-GAK-1 (1 μM), or dynasore (100 μM) for 2 hours. Surface IgM was detected using FITC-conjugated secondary antibody, and values were normalized to the 0-minute time point.

**S2D:** Representative flow cytometry histograms of surface BCR (FITC signal) at the 60-minute time point, corresponding to Fig. S2C.

**S2E:** Densitometric quantification of GAK protein levels following siGAK or NT siRNA transfection in DLBCL cells, normalized to B-actin and expressed as fold change relative to NT control.

**S2F:** Transferrin uptake assessed by flow cytometric MFI of CF®488A-transferrin in Ly1 cells pretreated with DMSO, SGC-GAK-1 (1 μM), or dynasore (100 μM). Values were normalized to DMSO control.

### Figure S3

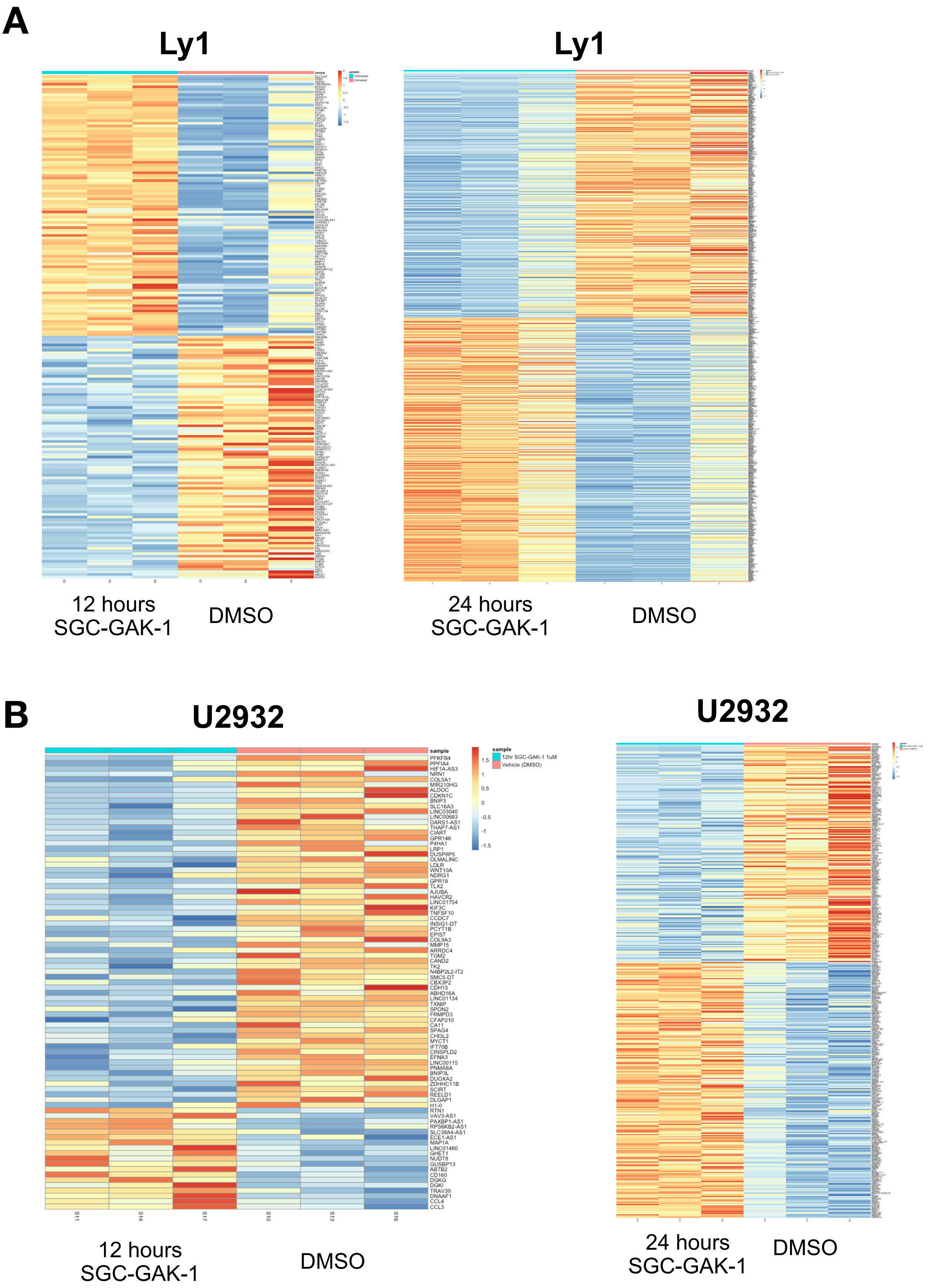
A
Ly1
Ly1
12 hours
SGC-GAK-1
DMSO
24 hours
SGC-GAK-1
DMSO
B
U2932
U2932
sample
12hr SGC-GAK-1 1uM
Vehicle (DMSO)
1.5
1
0.5
0
-0.5
-1
-1.5
PFKFB4
PPFIA4
HIF1A-AS3
NRN1
COL5A1
MIR210HG
ALDOC
CDKN1C
BNIP3
SLC16A3
LINC03040
LINC00683
DARS1-AS1
THAP7-AS1
CIART
GPR146
P4HA1
LRP1
DUSP8P5
OLMALINC
LDLR
WNT10A
NDRG1
GPR18
TLX2
AJUBA
HAVCR2
LINC01754
KIF3C
TNFSF10
CCDC7
INSIG1-DT
PCYT1B
EPIST
COL9A3
MMP15
ARRDC4
TGM2
CAND2
TK2
N4BP2L2-IT2
SMC5-DT
CBX3P2
CDH15
ABHD16A
LINC01134
TXNIP
SPON2
FRMPD3
CFAP210
CA11
SPAG4
CHI3L2
MYCT1
IFT70B
CRISPLD2
EFNA3
LINC00115
PNMA8A
BNIP3L
DUOXA2
ZDHHC11B
SCIRT
REELD1
DLGAP1
H1-0
RTN1
VAV3-AS1
PAXBP1-AS1
RPS6KB2-AS1
SLC38A4-AS1
ECE1-AS1
MAP1A
LINC01480
GHET1
NUDT8
GUSBP13
ABTB2
CD160
DGKG
DGKI
TRAV39
DNAAF1
CCL4
CCL3
S11
S14
S17
S10
S13
S16
12 hours
SGC-GAK-1
DMSO
24 hours
SGC-GAK-1
DMSO

### Figure S3

## C

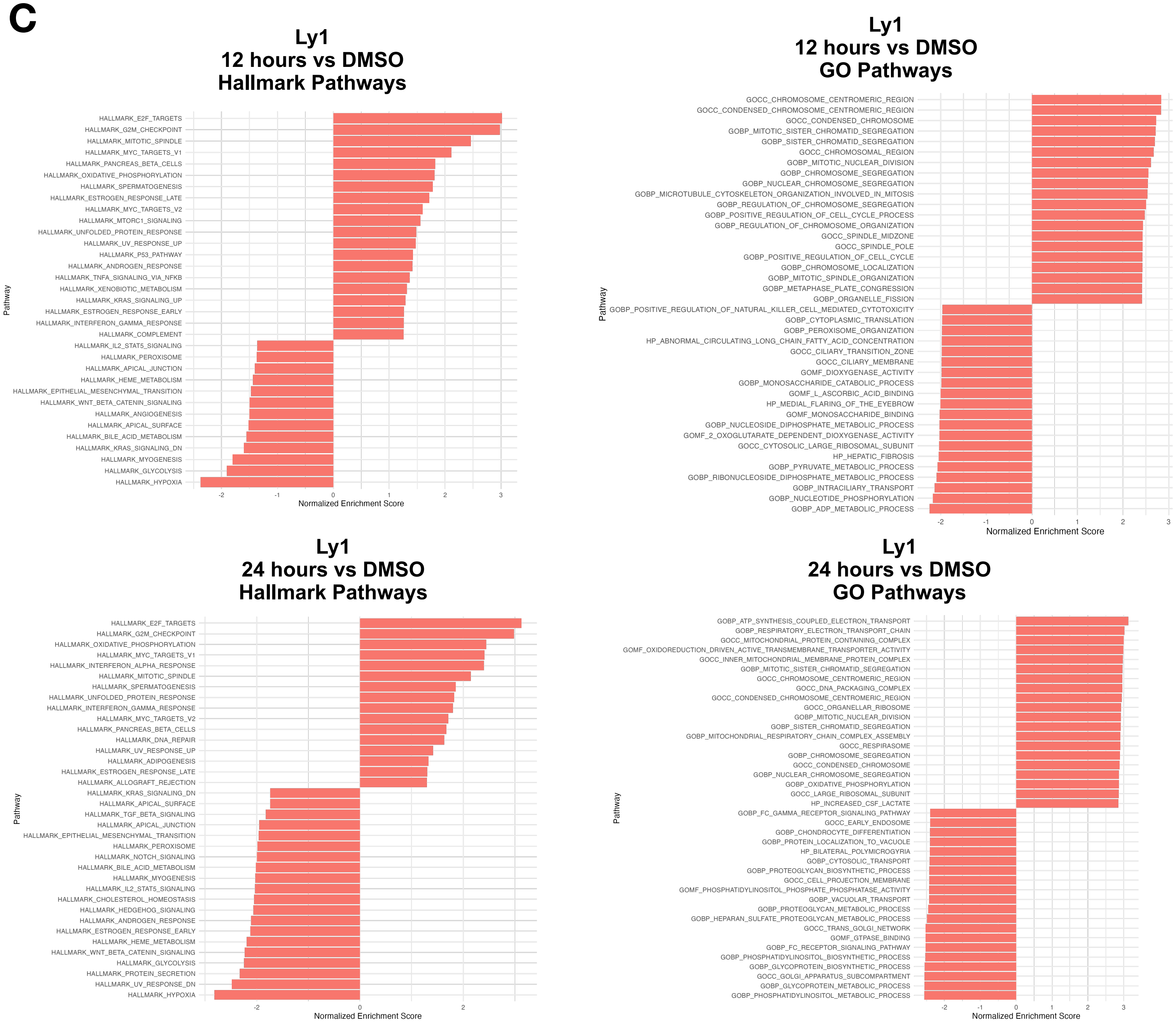
Ly1
12 hours vs DMSO
Hallmark Pathways
HALLMARK_E2F_TARGETS
HALLMARK_G2M_CHECKPOINT
HALLMARK_MITOTIC_SPINDLE
HALLMARK_MYC_TARGETS_V1
HALLMARK_PANCREAS_BETA_CELLS
HALLMARK_OXIDATIVE_PHOSPHORYLATION
HALLMARK_SPERMATOGENESIS
HALLMARK_ESTROGEN_RESPONSE_LATE
HALLMARK_MYC_TARGETS_V2
HALLMARK_MTORC1_SIGNALING
HALLMARK_UNFOLDED_PROTEIN_RESPONSE
HALLMARK_UV_RESPONSE_UP
HALLMARK_P53_PATHWAY
HALLMARK_ANDROGEN_RESPONSE
HALLMARK_TNFA_SIGNALING_VIA_NFKB
HALLMARK_XENOBIOTIC_METABOLISM
HALLMARK_KRAS_SIGNALING_UP
HALLMARK_ESTROGEN_RESPONSE_EARLY
HALLMARK_INTERFERON_GAMMA_RESPONSE
HALLMARK_COMPLEMENT
HALLMARK_IL2_STAT5_SIGNALING
HALLMARK_PEROXISOME
HALLMARK_APICAL_JUNCTION
HALLMARK_HEME_METABOLISM
HALLMARK_EPITHELIAL_MESENCHYMAL_TRANSITION
HALLMARK_WNT_BETA_CATENIN_SIGNALING
HALLMARK_ANGIOGENESIS
HALLMARK_APICAL_SURFACE
HALLMARK_BILE_ACID_METABOLISM
HALLMARK_KRAS_SIGNALING_DN
HALLMARK_MYOGENESIS
HALLMARK_GLYCOLYSIS
HALLMARK_HYPOXIA
Pathway
-2
-1
0
1
2
3
Normalized Enrichment Score
Ly1
12 hours vs DMSO
GO Pathways
GOCC_CHROMOSOME_CENTROMERIC_REGION
GOCC_CONDENSED_CHROMOSOME_CENTROMERIC_REGION
GOCC_CONDENSED_CHROMOSOME
GOBP_MITOTIC_SISTER_CHROMATID_SEGREGATION
GOBP_SISTER_CHROMATID_SEGREGATION
GOCC_CHROMOSOMAL_REGION
GOBP_MITOTIC_NUCLEAR_DIVISION
GOBP_CHROMOSOME_SEGREGATION
GOBP_NUCLEAR_CHROMOSOME_SEGREGATION
GOBP_MICROTUBULE_CYTOSKELETON_ORGANIZATION_INVOLVED_IN_MITOSIS
GOBP_REGULATION_OF_CHROMOSOME_SEGREGATION
GOBP_POSITIVE_REGULATION_OF_CELL_CYCLE_PROCESS
GOBP_REGULATION_OF_CHROMOSOME_ORGANIZATION
GOCC_SPINDLE_MIDZONE
GOCC_SPINDLE_POLE
GOBP_POSITIVE_REGULATION_OF_CELL_CYCLE
GOBP_CHROMOSOME_LOCALIZATION
GOBP_MITOTIC_SPINDLE_ORGANIZATION
GOBP_METAPHASE_PLATE_CONGRESSION
GOBP_ORGANELLE_FISSION
GOBP_POSITIVE_REGULATION_OF_NATURAL_KILLER_CELL_MEDIATED_CYTOTOXICITY
GOBP_CYTOPLASMIC_TRANSLATION
GOBP_PEROXISOME_ORGANIZATION
HP_ABNORMAL_CIRCULATING_LONG_CHAIN_FATTY_ACID_CONCENTRATION
GOCC_CILIARY_TRANSITION_ZONE
GOCC_CILIARY_MEMBRANE
GOMF_DIOXYGENASE_ACTIVITY
GOBP_MONOSACCHARIDE_CATABOLIC_PROCESS
GOMF_L_ASCORBIC_ACID_BINDING
HP_MEDIAL_FLARING_OF_THE_EYEBROW
GOMF_MONOSACCHARIDE_BINDING
GOBP_NUCLEOSIDE_DIPHOSPHATE_METABOLIC_PROCESS
GOMF_2_OXOGLUTARATE_DEPENDENT_DIOXYGENASE_ACTIVITY
GOCC_CYTOSOLIC_LARGE_RIBOSOMAL_SUBUNIT
HP_HEPATIC_FIBROSIS
GOBP_PYRUVATE_METABOLIC_PROCESS
GOBP_RIBONUCLEOSIDE_DIPHOSPHATE_METABOLIC_PROCESS
GOBP_INTRACILIARY_TRANSPORT
GOBP_NUCLEOTIDE_PHOSPHORYLATION
GOBP_ADP_METABOLIC_PROCESS
Pathway
-2
-1
0
1
2
3
Normalized Enrichment Score
Ly1
24 hours vs DMSO
Hallmark Pathways
HALLMARK_E2F_TARGETS
HALLMARK_G2M_CHECKPOINT
HALLMARK_OXIDATIVE_PHOSPHORYLATION
HALLMARK_MYC_TARGETS_V1
HALLMARK_INTERFERON_ALPHA_RESPONSE
HALLMARK_MITOTIC_SPINDLE
HALLMARK_SPERMATOGENESIS
HALLMARK_UNFOLDED_PROTEIN_RESPONSE
HALLMARK_INTERFERON_GAMMA_RESPONSE
HALLMARK_MYC_TARGETS_V2
HALLMARK_PANCREAS_BETA_CELLS
HALLMARK_DNA_REPAIR
HALLMARK_UV_RESPONSE_UP
HALLMARK_ADIPOGENESIS
HALLMARK_ESTROGEN_RESPONSE_LATE
HALLMARK_ALLOGRAFT_REJECTION
HALLMARK_KRAS_SIGNALING_DN
HALLMARK_APICAL_SURFACE
HALLMARK_TGF_BETA_SIGNALING
HALLMARK_APICAL_JUNCTION
HALLMARK_EPITHELIAL_MESENCHYMAL_TRANSITION
HALLMARK_PEROXISOME
HALLMARK_NOTCH_SIGNALING
HALLMARK_BILE_ACID_METABOLISM
HALLMARK_MYOGENESIS
HALLMARK_IL2_STAT5_SIGNALING
HALLMARK_CHOLESTEROL_HOMEOSTASIS
HALLMARK_HEDGEHOG_SIGNALING
HALLMARK_ANDROGEN_RESPONSE
HALLMARK_ESTROGEN_RESPONSE_EARLY
HALLMARK_HEME_METABOLISM
HALLMARK_WNT_BETA_CATENIN_SIGNALING
HALLMARK_GLYCOLYSIS
HALLMARK_PROTEIN_SECRETION
HALLMARK_UV_RESPONSE_DN
HALLMARK_HYPOXIA
Pathway
-2
0
2
Normalized Enrichment Score
Ly1
24 hours vs DMSO
GO Pathways
GOBP_ATP_SYNTHESIS_COUPLED_ELECTRON_TRANSPORT
GOBP_RESPIRATORY_ELECTRON_TRANSPORT_CHAIN
GOCC_MITOCHONDRIAL_PROTEIN_CONTAINING_COMPLEX
GOMF_OXIDOREDUCTION_DRIVEN_ACTIVE_TRANSMEMBRANE_TRANSPORTER_ACTIVITY
GOCC_INNER_MITOCHONDRIAL_MEMBRANE_PROTEIN_COMPLEX
GOBP_MITOTIC_SISTER_CHROMATID_SEGREGATION
GOCC_CHROMOSOME_CENTROMERIC_REGION
GOCC_DNA_PACKAGING_COMPLEX
GOCC_CONDENSED_CHROMOSOME_CENTROMERIC_REGION
GOCC_ORGANELLAR_RIBOSOME
GOBP_MITOTIC_NUCLEAR_DIVISION
GOBP_SISTER_CHROMATID_SEGREGATION
GOBP_MITOCHONDRIAL_RESPIRATORY_CHAIN_COMPLEX_ASSEMBLY
GOCC_RESPIRASOME
GOBP_CHROMOSOME_SEGREGATION
GOCC_CONDENSED_CHROMOSOME
GOBP_NUCLEAR_CHROMOSOME_SEGREGATION
GOBP_OXIDATIVE_PHOSPHORYLATION
GOCC_LARGE_RIBOSOMAL_SUBUNIT
HP_INCREASED_CSF_LACTATE
GOBP_FC_GAMMA_RECEPTOR_SIGNALING_PATHWAY
GOCC_EARLY_ENDOSOME
GOBP_CHONDROCYTE_DIFFERENTIATION
GOBP_PROTEIN_LOCALIZATION_TO_VACUOLE
HP_BILATERAL_POLYMICROGYRIA
GOBP_CYTOSOLIC_TRANSPORT
GOBP_PROTEOGLYCAN_BIOSYNTHETIC_PROCESS
GOCC_CELL_PROJECTION_MEMBRANE
GOMF_PHOSPHATIDYLINOSITOL_PHOSPHATE_PHOSPHATASE_ACTIVITY
GOBP_VACUOLAR_TRANSPORT
GOBP_PROTEOGLYCAN_METABOLIC_PROCESS
GOBP_HEPARAN_SULFATE_PROTEOGLYCAN_METABOLIC_PROCESS
GOCC_TRANS_GOLGI_NETWORK
GOMF_GTPASE_BINDING
GOBP_FC_RECEPTOR_SIGNALING_PATHWAY
GOBP_PHOSPHATIDYLINOSITOL_BIOSYNTHETIC_PROCESS
GOBP_GLYCOPROTEIN_BIOSYNTHETIC_PROCESS
GOCC_GOLGI_APPARATUS_SUBCOMPARTMENT
GOBP_GLYCOPROTEIN_METABOLIC_PROCESS
GOBP_PHOSPHATIDYLINOSITOL_METABOLIC_PROCESS
Pathway
-2
-1
0
1
2
3
Normalized Enrichment Score

### Figure S3

## D

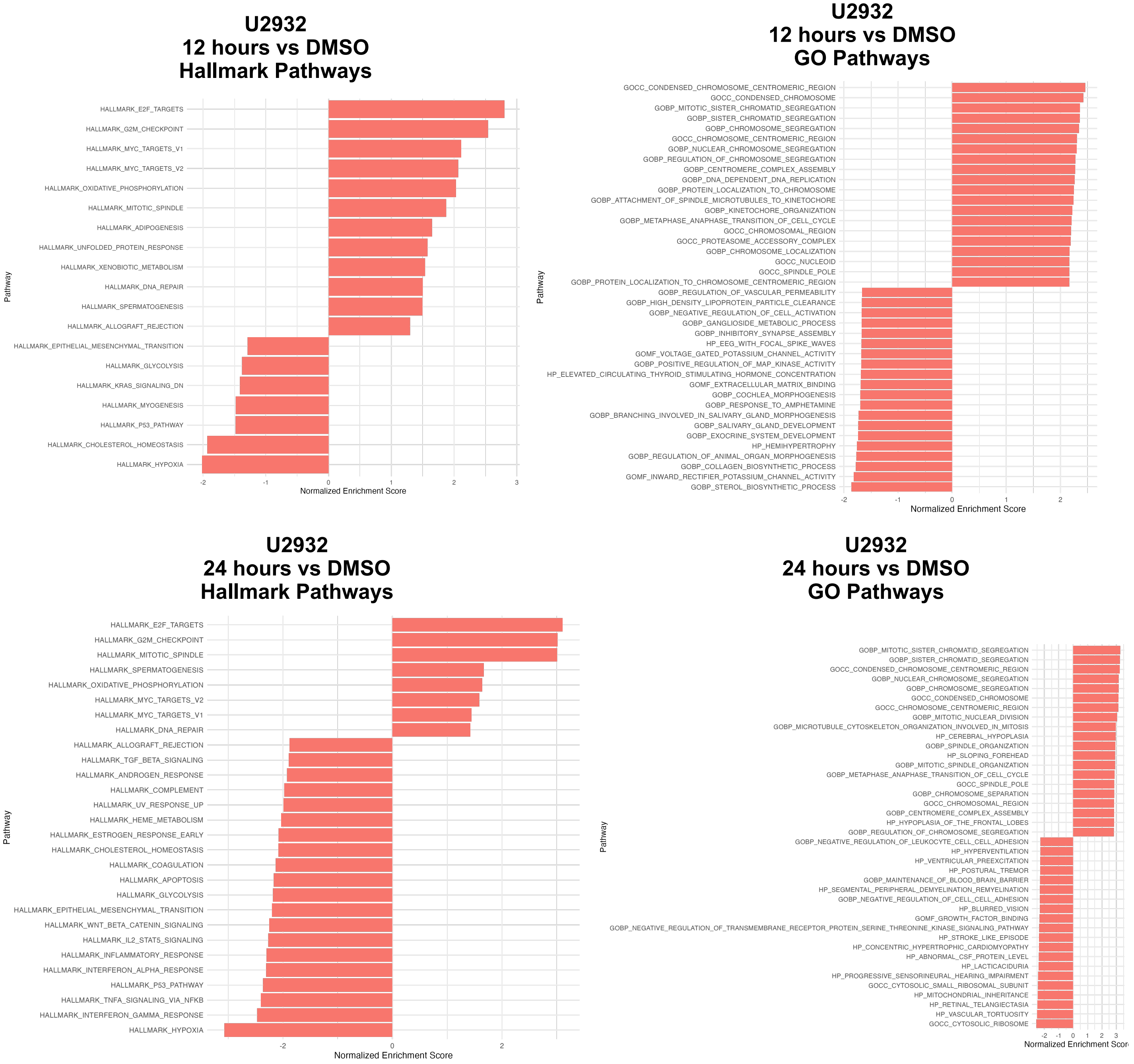

U2932
12 hours vs DMSO
Hallmark Pathways
HALLMARK_E2F_TARGETS
HALLMARK_G2M_CHECKPOINT
HALLMARK_MYC_TARGETS_V1
HALLMARK_MYC_TARGETS_V2
HALLMARK_OXIDATIVE_PHOSPHORYLATION
HALLMARK_MITOTIC_SPINDLE
HALLMARK_ADIPOGENESIS
HALLMARK_UNFOLDED_PROTEIN_RESPONSE
HALLMARK_XENOBIOTIC_METABOLISM
HALLMARK_DNA_REPAIR
HALLMARK_SPERMATOGENESIS
HALLMARK_ALLOGRAFT_REJECTION
HALLMARK_EPITHELIAL_MESENCHYMAL_TRANSITION
HALLMARK_GLYCOLYSIS
HALLMARK_KRAS_SIGNALING_DN
HALLMARK_MYOGENESIS
HALLMARK_P53_PATHWAY
HALLMARK_CHOLESTEROL_HOMEOSTASIS
HALLMARK_HYPOXIA
Pathway
-2
-1
0
1
2
3
Normalized Enrichment Score
U2932
12 hours vs DMSO
GO Pathways
GOCC_CONDENSED_CHROMOSOME_CENTROMERIC_REGION
GOCC_CONDENSED_CHROMOSOME
GOBP_MITOTIC_SISTER_CHROMATID_SEGREGATION
GOBP_SISTER_CHROMATID_SEGREGATION
GOBP_CHROMOSOME_SEGREGATION
GOCC_CHROMOSOME_CENTROMERIC_REGION
GOBP_NUCLEAR_CHROMOSOME_SEGREGATION
GOBP_REGULATION_OF_CHROMOSOME_SEGREGATION
GOBP_CENTROMERE_COMPLEX_ASSEMBLY
GOBP_DNA_DEPENDENT_DNA_REPLICATION
GOBP_PROTEIN_LOCALIZATION_TO_CHROMOSOME
GOBP_ATTACHMENT_OF_SPINDLE_MICROTUBULES_TO_KINETOCHORE
GOBP_KINETOCHORE_ORGANIZATION
GOBP_METAPHASE_ANAPHASE_TRANSITION_OF_CELL_CYCLE
GOCC_CHROMOSOMAL_REGION
GOCC_PROTEASOME_ACCESSORY_COMPLEX
GOBP_CHROMOSOME_LOCALIZATION
GOCC_NUCLEOID
GOCC_SPINDLE_POLE
GOBP_PROTEIN_LOCALIZATION_TO_CHROMOSOME_CENTROMERIC_REGION
GOBP_REGULATION_OF_VASCULAR_PERMEABILITY
GOBP_HIGH_DENSITY_LIPOPROTEIN_PARTICLE_CLEARANCE
GOBP_NEGATIVE_REGULATION_OF_CELL_ACTIVATION
GOBP_GANGLIOSIDE_METABOLIC_PROCESS
GOBP_INHIBITORY_SYNAPSE_ASSEMBLY
HP_EEG_WITH_FOCAL_SPIKE_WAVES
GOMF_VOLTAGE_GATED_POTASSIUM_CHANNEL_ACTIVITY
GOBP_POSITIVE_REGULATION_OF_MAP_KINASE_ACTIVITY
HP_ELEVATED_CIRCULATING_THYROID_STIMULATING_HORMONE_CONCENTRATION
GOMF_EXTRACELLULAR_MATRIX_BINDING
GOBP_COCHLEA_MORPHOGENESIS
GOBP_RESPONSE_TO_AMPHETAMINE
GOBP_BRANCHING_INVOLVED_IN_SALIVARY_GLAND_MORPHOGENESIS
GOBP_SALIVARY_GLAND_DEVELOPMENT
GOBP_EXOCRINE_SYSTEM_DEVELOPMENT
HP_HEMIHYPERTROPHY
GOBP_REGULATION_OF_ANIMAL_ORGAN_MORPHOGENESIS
GOBP_COLLAGEN_BIOSYNTHETIC_PROCESS
GOMF_INWARD_RECTIFIER_POTASSIUM_CHANNEL_ACTIVITY
GOBP_STEROL_BIOSYNTHETIC_PROCESS
Pathway
-2
-1
0
1
2
Normalized Enrichment Score
U2932
24 hours vs DMSO
Hallmark Pathways
HALLMARK_E2F_TARGETS
HALLMARK_G2M_CHECKPOINT
HALLMARK_MITOTIC_SPINDLE
HALLMARK_SPERMATOGENESIS
HALLMARK_OXIDATIVE_PHOSPHORYLATION
HALLMARK_MYC_TARGETS_V2
HALLMARK_MYC_TARGETS_V1
HALLMARK_DNA_REPAIR
HALLMARK_ALLOGRAFT_REJECTION
HALLMARK_TGF_BETA_SIGNALING
HALLMARK_ANDROGEN_RESPONSE
HALLMARK_COMPLEMENT
HALLMARK_UV_RESPONSE_UP
HALLMARK_HEME_METABOLISM
HALLMARK_ESTROGEN_RESPONSE_EARLY
HALLMARK_CHOLESTEROL_HOMEOSTASIS
HALLMARK_COAGULATION
HALLMARK_APOPTOSIS
HALLMARK_GLYCOLYSIS
HALLMARK_EPITHELIAL_MESENCHYMAL_TRANSITION
HALLMARK_WNT_BETA_CATENIN_SIGNALING
HALLMARK_IL2_STAT5_SIGNALING
HALLMARK_INFLAMMATORY_RESPONSE
HALLMARK_INTERFERON_ALPHA_RESPONSE
HALLMARK_P53_PATHWAY
HALLMARK_TNFA_SIGNALING_VIA_NFKB
HALLMARK_INTERFERON_GAMMA_RESPONSE
HALLMARK_HYPOXIA
Pathway
-2
0
2
Normalized Enrichment Score
U2932
24 hours vs DMSO
GO Pathways
GOBP_MITOTIC_SISTER_CHROMATID_SEGREGATION
GOBP_SISTER_CHROMATID_SEGREGATION
GOCC_CONDENSED_CHROMOSOME_CENTROMERIC_REGION
GOBP_NUCLEAR_CHROMOSOME_SEGREGATION
GOBP_CHROMOSOME_SEGREGATION
GOCC_CONDENSED_CHROMOSOME
GOCC_CHROMOSOME_CENTROMERIC_REGION
GOBP_MITOTIC_NUCLEAR_DIVISION
GOBP_MICROTUBULE_CYTOSKELETON_ORGANIZATION_INVOLVED_IN_MITOSIS
HP_CEREBRAL_HYPOPLASIA
GOBP_SPINDLE_ORGANIZATION
HP_SLOPING_FOREHEAD
GOBP_MITOTIC_SPINDLE_ORGANIZATION
GOBP_METAPHASE_ANAPHASE_TRANSITION_OF_CELL_CYCLE
GOCC_SPINDLE_POLE
GOBP_CHROMOSOME_SEPARATION
GOCC_CHROMOSOMAL_REGION
GOBP_CENTROMERE_COMPLEX_ASSEMBLY
HP_HYPOPLASIA_OF_THE_FRONTAL_LOBES
GOBP_REGULATION_OF_CHROMOSOME_SEGREGATION
GOBP_NEGATIVE_REGULATION_OF_LEUKOCYTE_CELL_CELL_ADHESION
HP_HYPERVENTILATION
HP_VENTRICULAR_PREEXCITATION
HP_POSTURAL_TREMOR
GOBP_MAINTENANCE_OF_BLOOD_BRAIN_BARRIER
HP_SEGMENTAL_PERIPHERAL_DEMYELINATION_REMYELINATION
GOBP_NEGATIVE_REGULATION_OF_CELL_CELL_ADHESION
HP_BLURRED_VISION
GOMF_GROWTH_FACTOR_BINDING
GOBP_NEGATIVE_REGULATION_OF_TRANSMEMBRANE_RECEPTOR_PROTEIN_SERINE_THREONINE_KINASE_SIGNALING_PATHWAY
HP_STROKE_LIKE_EPISODE
HP_CONCENTRIC_HYPERTROPHIC_CARDIOMYOPATHY
HP_ABNORMAL_CSF_PROTEIN_LEVEL
HP_LACTICACIDURIA
HP_PROGRESSIVE_SENSORINEURAL_HEARING_IMPAIRMENT
GOCC_CYTOSOLIC_SMALL_RIBOSOMAL_SUBUNIT
HP_MITOCHONDRIAL_INHERITANCE
HP_RETINAL_TELANGIECTASIA
HP_VASCULAR_TORTUOSITY
GOCC_CYTOSOLIC_RIBOSOME
Pathway
-2
-1
0
1
2
3
Normalized Enrichment Scor

### Figure S3

E

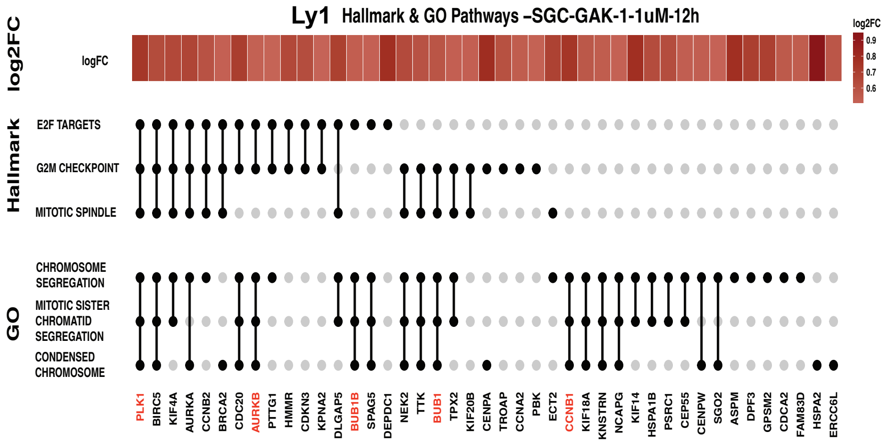


F

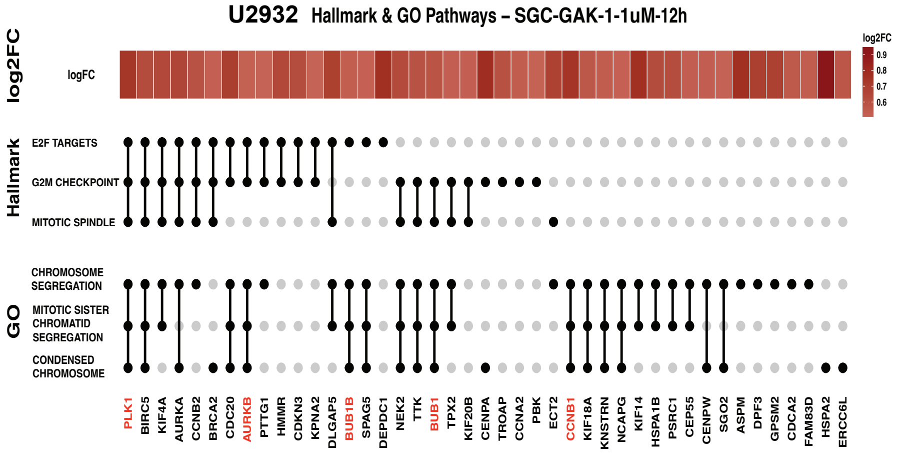


G

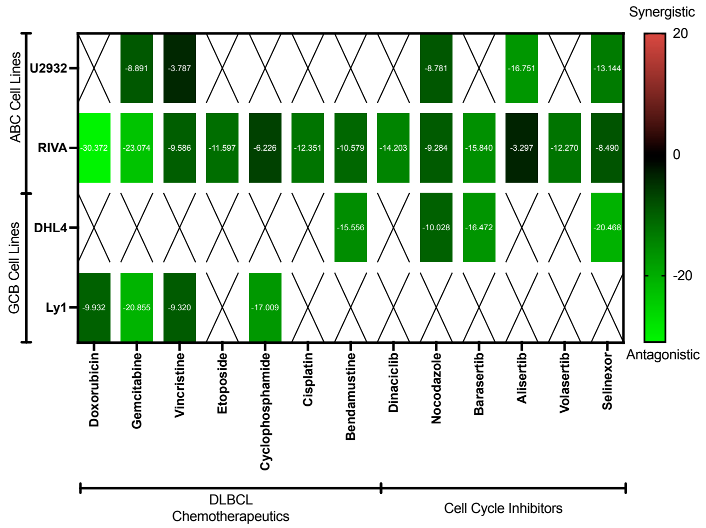

**Supplementary Figure S3** (previous pages)

**S3A-B:** Unsupervised clustering of differentially expressed genes following SGC-GAK-1 treatment (1 µM) for 12 and 24 hours relative to DMSO control ($p < 0.05$, $\log_2$ fold change > 0.5). (A) Ly1; (B) U2932.

**S3C-D:** FGSEA of Hallmark and Gene Ontology (GO) pathways in cells treated with SGC-GAK-1 (1 µM) for 12 and 24 hours relative to DMSO control. Enriched pathways derived from upregulated and downregulated genes (adjusted $p < 0.05$, $\log_2$ fold change > 0.5) are shown. (C) Ly1; (D) U2932.

**S3E-F:** Genes within enriched cell cycle–associated gene sets following SGC-GAK-1 treatment (1 µM) at 12 hours. Spindle assembly checkpoint (SAC) genes are highlighted in red. Gene sets are derived from pathways identified in Fig. 3D–E. (E) Ly1; (F) U2932.

**S3G:** Heatmap of Bliss synergy scores from U2932, RIVA, DHL4, and Ly1 cells treated with SGC-GAK-1 in combination with DLBCL chemotherapeutics and cell cycle inhibitors.

### Figure S4

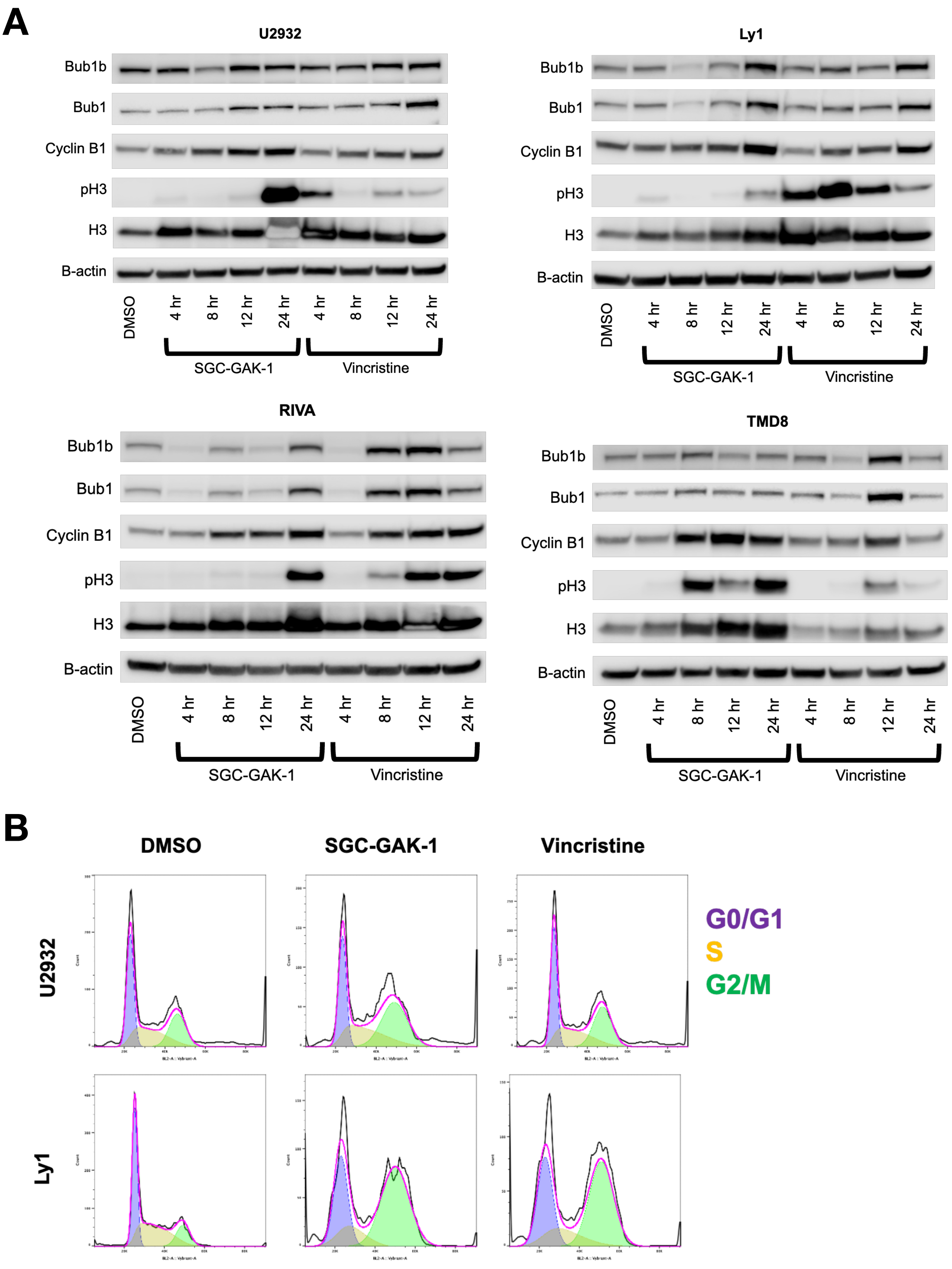
A
U2932
Ly1
RIVA
TMD8
Bub1b
Bub1
Cyclin B1
pH3
H3
B-actin
DMSO
4 hr
8 hr
12 hr
24 hr
SGC-GAK-1
Vincristine
B
DMSO
SGC-GAK-1
Vincristine
U2932
Ly1
G0/G1
S
G2/M

**Supplementary Figure S4** (previous page)

**S4A:** Immunoblot analysis of spindle assembly checkpoint and cell-cycle regulators in DLBCL cells treated with SGC-GAK-1 (1 μM), Vincristine (1 nM), or DMSO control over a time course.

**S4B:** Representative flow cytometry histograms of DNA content (Vybrant DyeCycle Orange) used for cell cycle analysis in U2932 and Ly1 cells treated with SGC-GAK-1 (1 μM), vincristine (1 nM), or DMSO for 24 hours, corresponding to Fig. 4B.

#### Figure S5

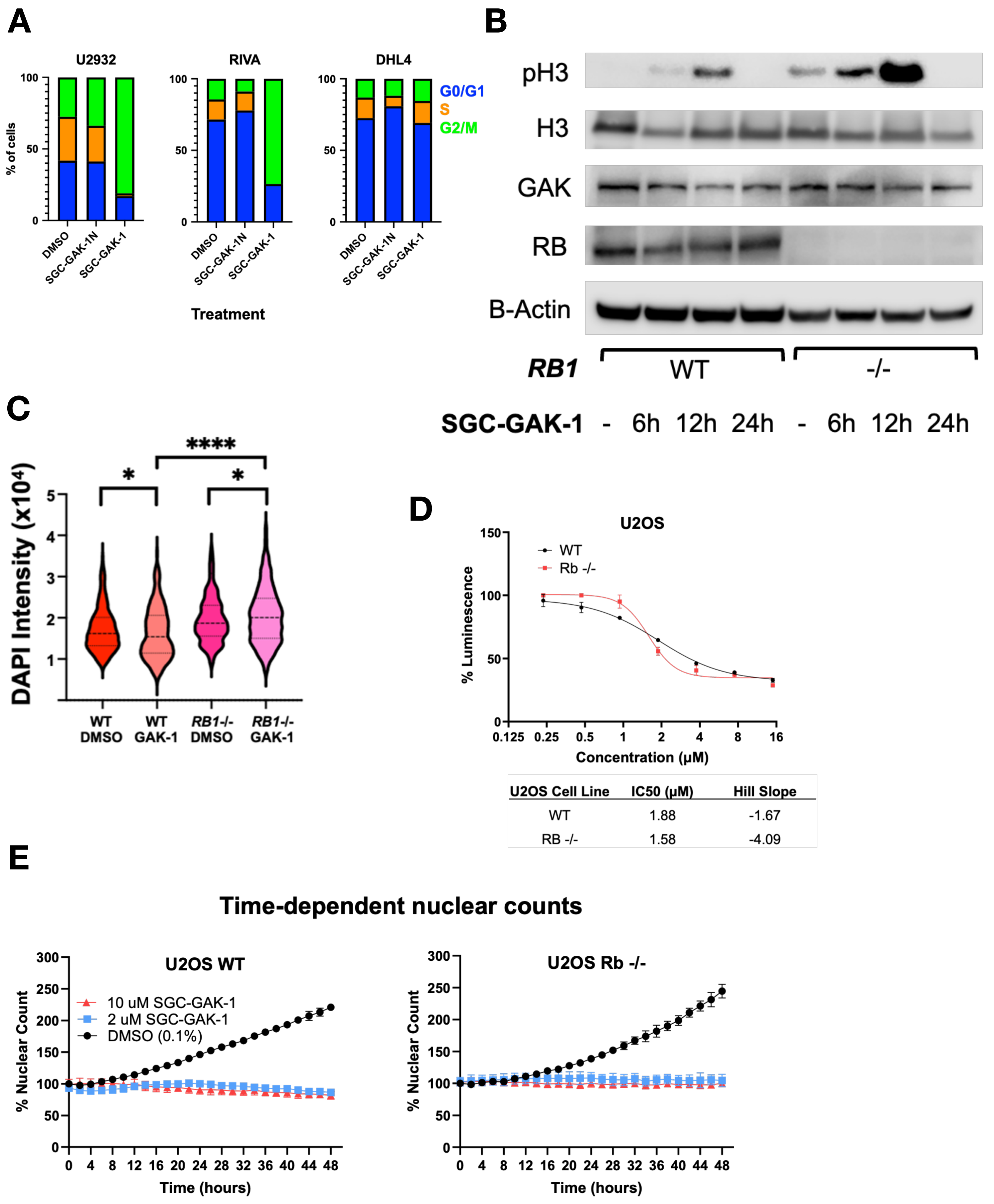


| U2OS Cell Line | IC50 (μM) | Hill Slope |
|---|---|---|
| WT | 1.88 | -1.67 |
| RB -/- | 1.58 | -4.09 |

**Supplementary Figure S5** (previous page)

**S5A:** Cell cycle distribution determined by flow cytometry (DNA content analysis) in DLBCL cells treated with SGC-GAK-1 (10 μM), SGC-GAK-1N (10 μM), or DMSO for 24 hours, corresponding to Fig. 5D.

**S5B:** Immunoblot analysis of RB, GAK, phospho-histone H3 (pH3), and total H3 in U2OS WT and $RB^{-}/^{-}$ cells following time-course treatment with SGC-GAK-1 (1 μM).

**S5C:** Violin plots quantifying chromosomal condensation via DAPI staining in WT and RB1–/– U2OS cells treated with DMSO or SGC-GAK-1 (1 μM, 24 hr), corresponding to Fig. 5E (****p < 0.0001, *p < 0.05).

**S5D:** Dose–response cell viability assay of U2OS WT and RB-/- cells treated with SGC-GAK-1 for 48 hrs, assessed by CellTiter-Glo (CTG).

**S5E:** Time-course of percent nuclear count in U2OS WT and $RB^{-}/^{-}$ cells treated with SGC-GAK-1 (2 μM or 10 μM) or DMSO, measured by live-cell imaging and normalized to the 0-hour time point.

Figure S6

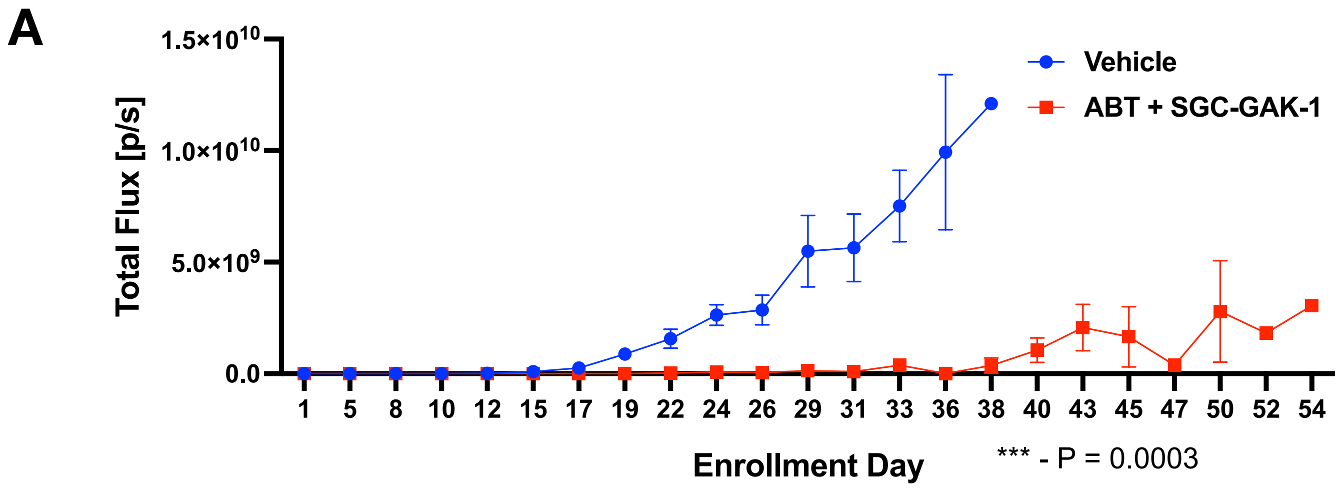
A
1.5×10^10
1.0×10^10
5.0×10^9
0.0
Total Flux [p/s]
Vehicle
ABT + SGC-GAK-1
1
5
8
10
12
15
17
19
22
24
26
29
31
33
36
38
40
43
45
47
50
52
54
Enrollment Day
*** - P = 0.0003

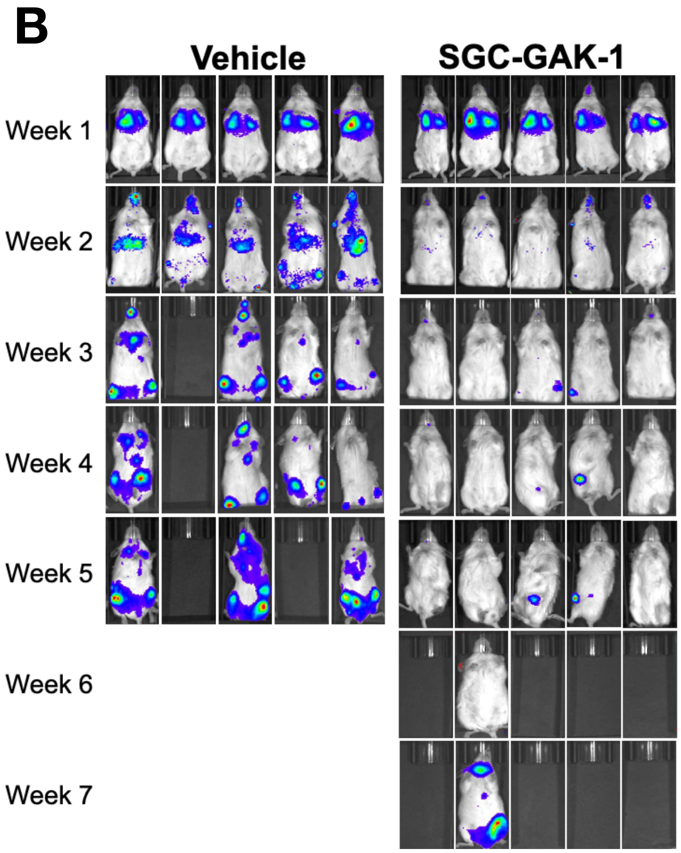
B
Vehicle
SGC-GAK-1
Week 1
Week 2
Week 3
Week 4
Week 5
Week 6
Week 7

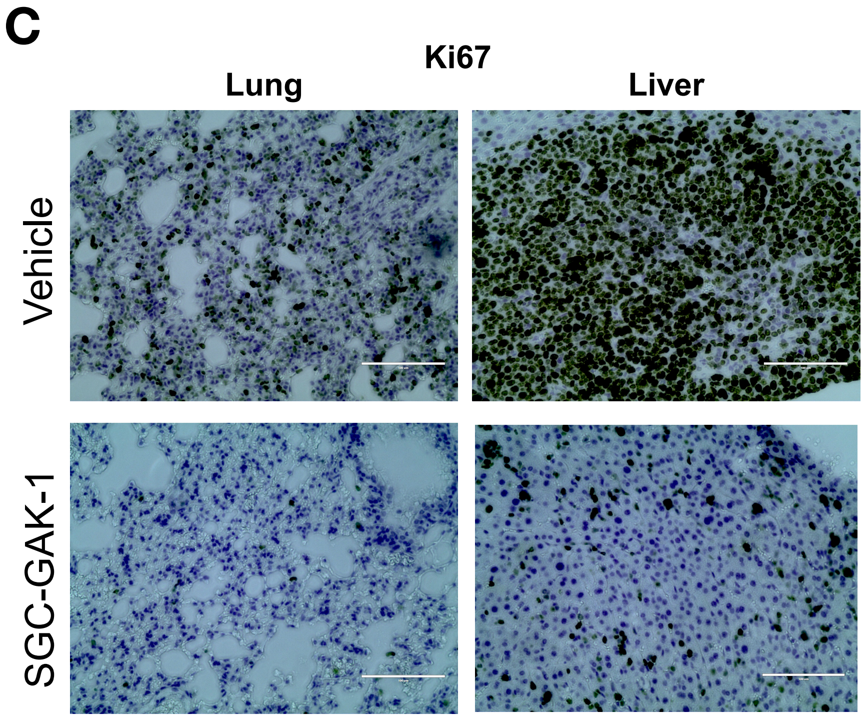
C
Ki67
Lung
Liver
Vehicle
SGC-GAK-1

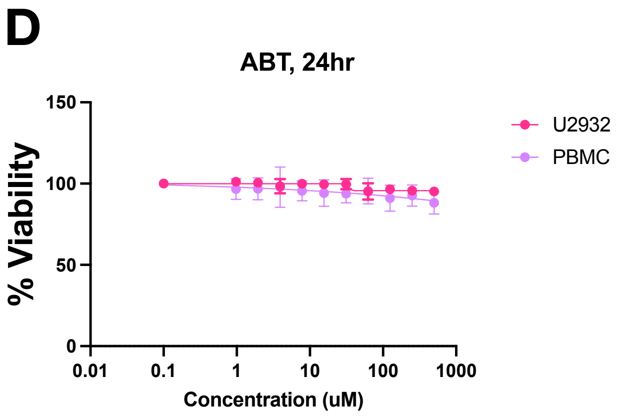
D
ABT, 24hr
% Viability
150
100
50
0
0.01
0.1
1
10
100
1000
Concentration (uM)
U2932
PBMC

**Supplementary Figure S6** (previous page)

**S6A:** Tumor burden quantified as average signal intensity (± SEM) by IVIS imaging in NSG mice bearing luciferase-expressing U2932 xenografts treated with SGC-GAK-1 + ABT or vehicle control. *p = 0.0003 (two-way ANOVA).

**S6B:** Representative IVIS images of NSG mice bearing luciferase-expressing U2932 xenografts treated with SGC-GAK-1 + ABT or vehicle control.

**S6C:** Immunohistochemical (IHC) staining for Ki67 in tumor tissues from lung and liver of NSG mice treated with SGC-GAK-1 + ABT or vehicle control.

**S6D:** Dose–response cell viability of U2932 cells and PBMCs following ABT treatment for 24 hours.
